# The molecular architecture of tunneling nanotubes

**DOI:** 10.64898/2026.05.27.728322

**Authors:** Eva P Karasmanis, Siyu Chen, Farhaz Shaikh, Robert G. Abrisch, Alex Flaherty, Rafael A. Badell-Grau, Margot Riggi, Landon Vu Nguyen, Joshua Hutchings, Tamar Basiashvili, Nicholas Lattal, Stephanie Cherqui, Elizabeth Villa, Samara L Reck-Peterson

## Abstract

Tunneling nanotubes (TNTs) are thin intercellular bridges that mediate the exchange of proteins, organelles, and nucleic acids between neighboring cells. They are enriched in tumor cells, implicated in chemotherapy resistance, induced in models of aggregation-based diseases, and their formation is stimulated by viruses that use TNTs to enhance infection. Despite their broad relevance and therapeutic potential, TNT morphology and function remain poorly understood, owing to the absence of clear morphological criteria and the limitations of light microscopy. Here, we establish two complementary systems to study TNT formation and function: stimulation with the pseudorabies viral kinase US3 to model viral transmission, and treatment of acute monocytic leukemia THP-1 cells with daunorubicin to model chemotherapy resistance. Using live-cell imaging, we characterize cytoskeletal organization and bidirectional lysosome transport in both contexts, and apply cryo-correlative light and electron microscopy (cryo-CLEM) with cryogenic electron tomography (cryo-ET) to visualize TNTs in their native state at molecular resolution. We show that TNTs display a rich molecular architecture, comprising actin filaments, microtubules, intermediate filaments, active ribosomes, and diverse organelles including multivesicular bodies, autophagosomes, and lysosomes. Sub-nanometer microtubule reconstructions reveal mixed polarity within individual TNTs, suggesting that both connected cells actively contribute to TNT formation and cargo trafficking. This organization is conserved across both systems, implying that TNT biogenesis reflects a shared cellular program rather than a context-specific response. Our findings provide the first structural framework for TNTs, revealing an unexpectedly rich molecular architecture and laying the groundwork for understanding how TNTs orchestrate intercellular communication in disease.

## Introduction

Cell-to-cell communication is essential to multicellularity, and cells have evolved a diverse repertoire of mechanisms to exchange cell signals and materials with their neighbors. Amongst the most recently described mechanisms of such exchange are tunneling nanotubes (TNTs), which are thin, long, membrane suspended bridges formed between two cells, providing a direct cytoplasmic continuum for the exchange of cell materials including organelles, proteins, ions, nucleic acids and pathogens. First described in 2004 as “nanotubular highways” mediating organelle transport in cultured cells^1,2^, TNTs have since been observed in an expanding range of cell types and physiological and pathological contexts *in vivo* and in culture.

Structurally, TNTs are defined by their elevation above the substrate, distinguishing them from filopodia, and by their cytoplasmic continuity with the cells they connect. They typically range from 50 nm to over 1 μm in diameter and can span tens to hundreds of micrometers. Recent ultrastructure studies suggest that TNTs are actin-based structures^1,3–5^, while other studies suggest TNTs may also contain microtubules^2,6–10^. Notably, organelle and vesicular transport through TNTs has been reported in many cell types, and the recorded motile behaviors are consistent with active microtubule-based transport^1,11–15^. However, whether TNTs are truly open-ended structures permitting bidirectional cargo exchange, or closed, remains a source of ongoing debate^16–19^. Distinguishing bona fide TNTs from morphologically related structures (e.g. filopodia, retraction fibers, cytonemes, or lingering intercellular bridges) is further complicated by the absence of definitive molecular markers, limited structural data, and the likelihood that multiple classes of thin intercellular connections coexist *in vivo* with overlapping functions^15,18–20^.

Notwithstanding these challenges, the functional relevance of TNTs across diverse biological contexts is increasingly compelling. In the nervous system, TNTs have emerged as critical conduits for the intercellular spread of pathological cargo in neurodegeneration, including prion-like aggregates, α-synuclein, tau, and mutant huntingtin^7,10,21–24^. Microglia have been shown to connect with neurons and one another through TNTs to collectively manage the burden of protein aggregates, distributing them from burdened cells to naïve neighbors for degradation while simultaneously returning healthy mitochondria in a cellular model of Parkinson’s disease ^25,26^. TNTs are also being explored as an important avenue for therapeutics. This interest is driven in part by the finding that a stem cell gene therapy for the lysosomal storage disorder cystinosis, is mediated through a TNT-dependent mechanism^27–29^.

Evidence for TNT-mediated communication is particularly compelling in cancer and viral infection. *In vivo* imaging of tumor models has identified extensive networks of long, thin protrusions (also termed tumor microtubes) that interconnect cancer cells and support calcium signaling, mitochondrial transfer, and resistance to chemotherapy^30–38^. Numerous viruses, including HIV, SARS-CoV-2, and herpesviruses, have been shown to induce and exploit TNT formation for cell-to-cell spread, accessing cell types that lack conventional entry receptors and evading neutralizing antibodies^11,39–44^.

While TNTs have been most extensively studied in the context of disease, evidence is mounting that they also serve fundamental roles in normal physiology. In the immune system, membrane nanotubes have been visualized connecting dendritic cells, macrophages, and T cells within lymphoid tissues, where they facilitate long-range calcium flux and antigen-dependent signaling beyond direct cell–cell contact^43,45^. Recent *in vivo* studies have further identified TNT-like structures capable of long-range intercellular cargo transfer in developing zebrafish embryos and in the mouse neonatal cerebellum, cardiac, and neural systems, underscoring their broader relevance in development, organogenesis, and tissue homeostasis^20,46–48^.

Despite this growing body of evidence, TNTs remain controversial. Their nanoscale diameter, dynamic nature, sensitivity to conventional fixation, and lack of distinctive molecular markers have made rigorous structural and functional characterization challenging. The structural underpinnings of TNT biology are poorly defined. Here, we systematically survey primary and cultured cells to establish TNT occurrence and leverage two complementary model systems that induce TNT formation: cells treated with the herpesvirus kinase US3 and daunorubicin-treated pediatric leukemia THP-1 cells. We quantitatively analyze TNT length, cytoskeletal content and polarity, and bidirectional lysosomal transport. We use cryogenic correlative light electron microscopy (CLEM) combined with cryogenic electron tomography (cryo-ET) to examine the TNT molecular landscape. We find that TNTs exhibit a complex and highly organized architecture where multiple cytoskeletal systems (actin, microtubules, and intermediate filaments) co-exist with a dense, dynamic and translationally active intracellular environment enriched with vesicles and organelles, including lysosomes, multivesicular bodies, mitochondria, autophagosomes and endoplasmic reticulum. Notably, although TNTs are likely initially actin based, they are permeated by dynamic, opposite-polarity microtubules, suggesting that cells on either end of a TNT contribute the formation of the intercellular highways that can rapidly and bidirectionally exchange large cellular objects. We show that this TNT molecular organization is conserved between different cell lines and physiological contexts, suggesting TNT formation is a robust mechanism for intercellular communication to mitigate cytotoxic stress. Taken together, our results reveal the unexpectedly rich and conserved molecular architecture of TNTs across diverse biological contexts.

## Results

### TNTs form spontaneously across diverse cell types

To systematically assess TNT formation, we surveyed a panel of primary and immortalized cell lines, including primary mouse embryonic fibroblasts (MEFs), U2OS, HeLa, and A549 cells. Filamentous actin and microtubules were identified by immunofluorescence (Fig. 1a). TNTs were defined as thin, actin-positive membranous connections between two cells, longer than 4 μm. Mitotic cells, including those in late cytokinesis, were excluded from analysis. For each TNT, length and the presence or absence of tubulin staining were recorded. Across all cell types, TNTs formed spontaneously at a consistent basal frequency of ∼5%-10% (Fig. 1a), indicating that TNT biogenesis is a general feature of diverse cell types rather than a cell line-specific phenomenon. To assess intercellular transport through TNTs, we focused on primary cells. We generated stable mouse fibroblast lines via lentiviral delivery of the lysosomal protein cystinosin (CTNS) tagged with green fluorescent protein (GFP). Using live-cell imaging we visualized bidirectional lysosomal transport through TNTs, confirming active intercellular exchange of lysosomes (Fig. 1b; Supplementary Movie 1)^29^. TNT length varied widely, ranging from a few microns to several hundred microns, consistent with previous reports^1,43,49–52^ (Fig. 1c).

**Figure 1.**
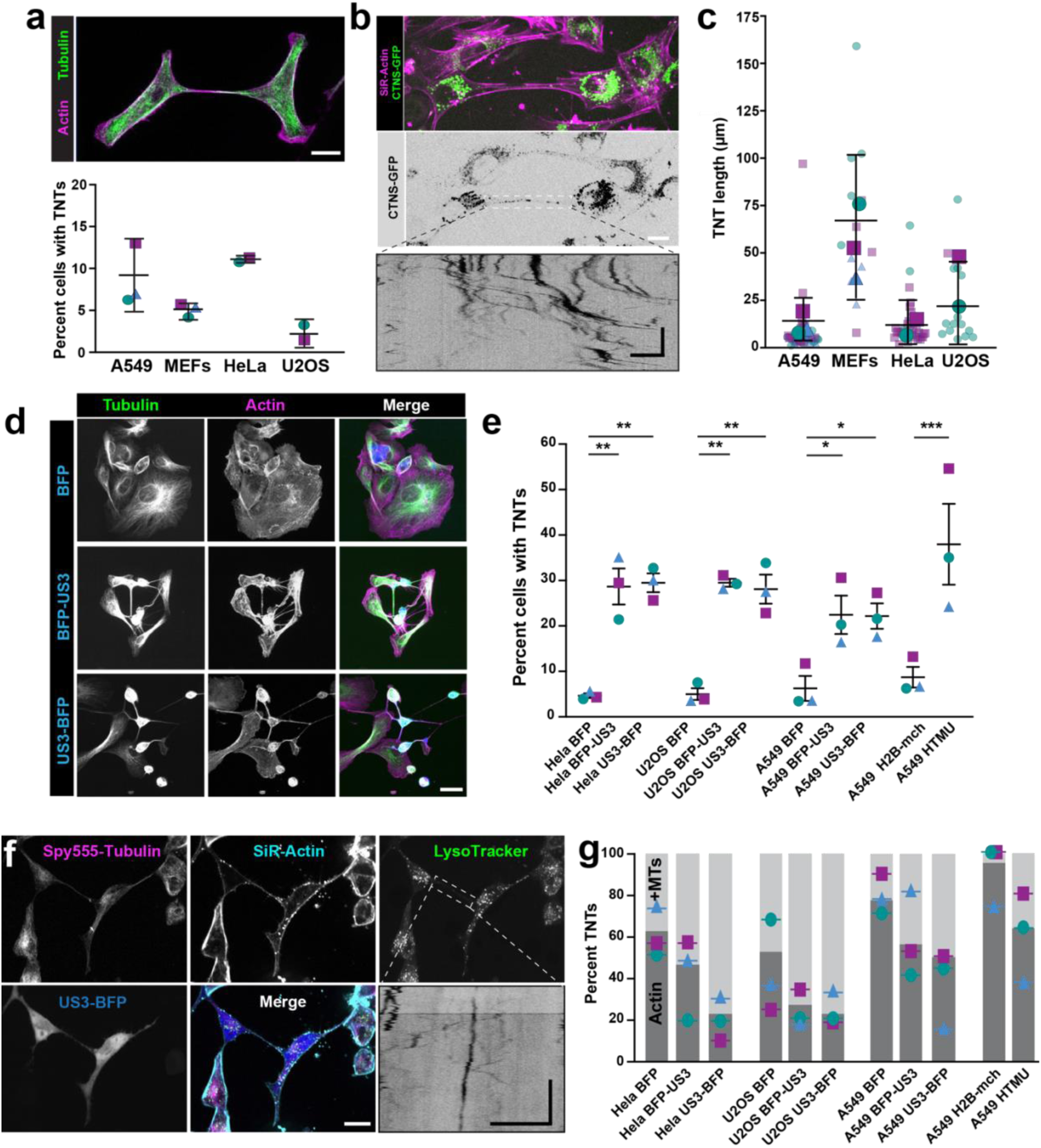
TNTs form in diverse cell types and US3 kinase promotes robust TNT formation. **a.** Top: Representative image of A549 cells with a TNT. Cells were stained for filamentous actin (phalloidin 647) and microtubules (DM1a; Sigma). Bottom: Basal TNT formation across primary mouse embryonic fibroblasts (MEFs) and immortalized cell lines (A549, HeLa, U2OS). Across cell lines, the frequency of TNT formation was ∼5-10%. Different shapes and colors denote different biological replicates. Total cells analyzed and corresponding TNT counts for each replicate are as follows: A549: n = 96/166/99 cells and 6/22/6 TNTs, MEFs: n = 131/71/81 cells and 5/4/4 TNTs, U2OS: n = 169/75 cells, 6/1 TNTs, HeLa: n = 72/254 cells and 8/27 TNTs. **b.** Representative image of MEFs stably expressing cystinosin (CTNS-GFP). SiR-actin was used to visualize filamentous actin and TNTs. Kymograph (bottom) shows bidirectional lysosomal transport through the TNT shown above. **c.** Quantification of TNT length in A549, primary MEFs, HeLa and U2OS cells. Superplots show all individual data points for each of the two or three biological replicates with different shapes and colors correspond to different biological replicates. Larger, opaque shapes denote the mean of each biological replicate. Bar graph and error bars indicate the mean ± SEM. A549: n = 6/22/6 TNTs, MEFs: n = 5/4/4 TNTs, U2OS: n = 6/1 TNTs, HeLa: n = 8/27 TNTs. **d.** Representative images of U2OS cells expressing BFP (control) or pseudorabies viral kinase US3 fused to BFP at the N- (BFP-US3) or C-terminus (US3-BFP). Cells were fixed and stained for filamentous actin (phalloidin 647) and microtubules (anti-mouse α-tubulin 568). **e.** Quantification of the percent cells with TNTs in HeLa, U2OS, and A549 cells expressing the US3 kinase. Three biological replicates were performed. Bars represent mean ± SEM. Different shapes and colors denote individual replicates where n > 200 cells per replicate. HTMU; tandem lentiviral construct expressing H2B-mCherry and untagged US3 kinase. Statistical analysis was done using a one-way ANOVA with Tukey’s multiple comparison test. HeLa BFP and HeLa BFP-US3 ** P = 0.0052, HeLa BFP and HeLa US3-BFP ** P = 0.0037, U2OS BFP and U2OS BFP-US3 ** P = 0.0042, U2OS BFP and U2OS US3-BFP ** P = 0.0049, A549 BFP and A549 BFP-US3 * P = 0.0318, A549 BFP and A549 US3-BFP * P =0.0341, A549 H2B-mCherry and A549 HTMU ***P = 0.0005. **f.** Representative image of TNTs formed in HeLa cells expressing US3-BFP. Lysosomes (LysoTracker Green), microtubules (SPY555-tubulin), and filamentous actin (SiR-actin) were imaged in live cells. Kymograph shows the bidirectional transport of lysosomes within the outlined TNT. **g.** Stacked bar graph shows the percent TNTs positive for actin only (dark gray), or for both actin and microtubules (light gray). Shapes and colors denote the percent actin-only containing TNTs in three biological replicates. HeLa BFP: n = 3/2/1 actin TNTs, n = 0/4/4 +MTs TNTs, HeLa BFP-US3: n = 28/22/15 actin TNTs, n = 7/16/22 +MTs TNTs, HeLa US3-BFP: n = 62/24/25 actin TNTs, n = 15/7/11 +MTs TNTs. U2OS BFP: n = 18/5/8 actin TNTs, n = 18/2/6 +MTs TNTs, U2OS BFP-US3: n = 30/31/21 actin TNTs, n = 8/16/7 +MTs TNTs, U2OS US3-BFP: n=34/22/38 actin TNTs, n = 9/8/11 +MTs TNTs. A549 BFP: n = 2/3/5 actin TNTs, n=5/11/11 +MTs TNTs, A549 BFP-US3: n = 31/15/16 actin TNTs, n = 14/18/20 +MTs TNTs, A549 US3-BFP: n=26/21/11 actin TNTs, n = 23/27/15 +MTs TNTs. A549 H2B-mcherry: n =11/10/6 actin TNTs n = 0/2/0 +MTs TNTs, A549 HTMU: n = 12/26/11 actin TNTs, n = 28/41/16 +MTs TNTs TNTs quantified from the >200 cells per replicate in e. MT; microtubule. Scale bars: 20 μm in all panels except for kymographs; kymographs in b and f: horizontal 10 μm, vertical 5 min.

### US3 kinase expression drives robust TNT induction across diverse cell lines

Because TNTs formed relatively infrequently under these conditions, to achieve robust, reproducible TNT induction, we leveraged the pseudorabies viral serine/threonine kinase US3, which is conserved across the Alphaherpesvirinae and a well-characterized driver of TNT formation^44,53^. Expression of US3 alone was sufficient to dramatically increase the proportion of TNT-positive cells to ∼35% across multiple lines (Fig. 1d, Extended Data Fig. 1b) generating stable, microtubule-containing TNTs that have previously been shown to support viral spread even in the presence of neutralizing antibodies^53^. Tagging US3 kinase with blue fluorescent protein (BFP) at either the N- or C-terminus did not alter its ability to induce TNT formation (Fig. 1d-e). In A549 cells, which exhibit low transfection efficiency, lentiviral delivery of US3 achieved comparable levels of TNT induction (Fig. 1d). Live-cell imaging of US3-BFP expressing cells stained with LysoTracker, SPY555-tubulin and SiR-actin, showed that bidirectional lysosomal transport occurred in US3 kinase-induced TNTs, suggesting they are functionally comparable to endogenous TNTs (Fig. 1f; Supplementary Movie 2). A larger fraction of US3-induced TNTs contained microtubules, whereas TNTs in control conditions (non US3-induced) were predominantly actin-only (Fig. 1g). Lysosomal transport was observed in TNTs that were positive for microtubules and lysosome velocities were consistent with microtubule-based transport (Fig. 1b and 1f). TNTs containing actin alone had an average length of ∼10 μm across all cell lines, whereas TNTs containing both actin and microtubules were generally longer (∼35 μm). The average TNT length did not change upon induction by US3 (Extended Data Fig. 1b). Together, these results establish US3 kinase expression as a reliable and versatile strategy for TNT induction across diverse cell backgrounds.

### Thin, actin-only TNTs are closed or unresolved at their endpoints

To directly visualize the molecular composition and architecture of TNTs and address the ongoing debate regarding their existence as bona fide modes of intercellular communication, we performed *in situ* CLEM and cryo-ET ^54,55^. We imaged TNTs formed in A549 and HeLa cells expressing US3 kinase (via lentivirus or US3-BFP, respectively) (Extended Data Fig. 2 and 3). On the lower end of TNT thickness (∼100 nm) (Extended Data Fig. 4a), we observed intercellular protrusions that were closed; a bilayer was present somewhere along the tubule, rendering the interior non-contiguous between cells, or whose endpoint continuity could not be confidently resolved (Extended Data Fig. 2a–c). These structures contained actin filaments, a few vesicles (Extended Data Fig. 2a), and no microtubules. This is consistent with prior cryo-ET studies of TNTs^3,42^, which characterized thin actin-containing structures, therefore we categorized these structures separately (Extended Data Fig. 2 a-c) and focused our analysis on open, microtubule-containing TNTs.

### Cryo-ET reveals a crowded and diverse organelle landscape within TNTs

Only intercellular bridges whose end-to-end openness could be confidently resolved in medium-magnification overview montages were taken forward as bona fide open TNTs for tomogram acquisition (Fig. 2 a-b; Extended Data Fig. 2 d-e; Extended Data Fig. 3). We observed long, membrane-enclosed intercellular bridges spanning several microns between distant cells (Fig. 2a; Extended Data Fig. 2d; Extended Data Fig. 3), which were thicker compared to the thin, actin structures reported previously^3,42^(Extended Data Fig. 4a). CLEM confirmed that these structures were positive for lysosomes (LysoTracker), actin (SiR-actin), and microtubules (SPY555-tubulin) (Fig. 2a). Overview montages and tomograms spanning the TNT length confirmed that these TNTs were open, and had a continuous lumen connecting the two cells, with no observable membrane barriers (Fig. 2b-c; Extended Data Fig. 2d-e; Extended Data Fig. 3). Our data revealed that TNTs contain a rich and diverse vesicular population. Many vesicles were large (>150 nm in diameter) (Extended Data Fig. 4c), consistent with late endolysosomal vesicles. Using CLEM-ET we were able to identify autophagosomes, mitochondria, multivesicular bodies and lysosomes. Smaller vesicles (<150 nm) were also abundant, suggesting the presence of endosomes and signaling vesicles (Extended Data Fig. 4c).

**Figure 2.**
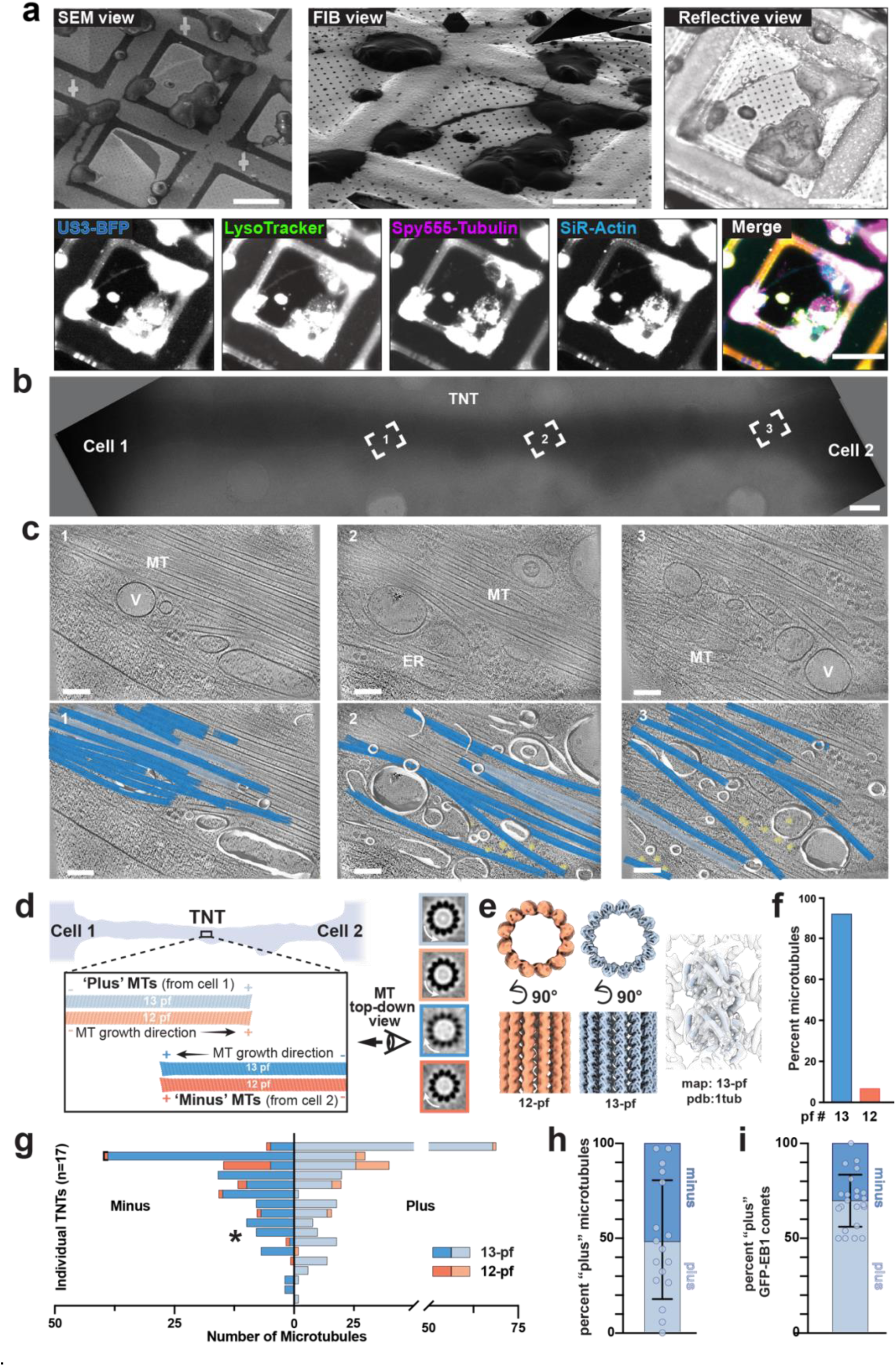
In-situ cryo–ET reveals the diverse organelle landscape within TNTs. **a.** Scanning electron microscopy and focused-ion-beam views of TNTs on grids. Reflective microscopy followed by cryo-widefield fluorescent imaging identified US3-BFP-induced TNTs positive for lysosomes, actin and microtubules. **b.** Cryo-TEM montage showing a representative TNT connecting two distant cells. White dashed boxes (1-3) mark representative positions of tilt–series acquisition along the nanotube where microtubules were used for subtomogram averaging. Higher zoom of this montage and the full panel of tomogram acquisition are in Extended Data Fig. 3a. **c.** Enlarged tomographic slices (10 nm thick) (1, 2, 3 in (b)) highlight the crowded interior of the TNT. Labeled features include microtubules (MT), endoplasmic reticulum (ER), and vesicles (V). Bottom: segmentation and spatial reconstruction of the regions shown in (b). All microtubules in this tomogram are 13-pf. Vesicular and ER membranes (white) and ribosomes (yellow) are rendered to illustrate their 3D spatial organization within the TNT. **d.** Illustration showing our microtubule polarity determination workflow. Microtubules are initially picked uni-directionally. Representative cross-sections from per-microtubule alignment reveal the characteristic skewing direction, allowing the assignment of individual microtubules as ‘Plus’ or ‘Minus’, corresponding to origin from cell1 and cell2, respectively. Microtubules are color-coded consistently across all figures by their determined polarity and pf-architecture: blue tones represent 13-pf, while orange tones represent 12-pf. Within each architecture, light shades indicate ‘Plus’ directed microtubules, and dark shades indicate ‘Minus’ directed microtubules. **e.** 3D reconstructions of 12-protofilament (12-pf, orange) and 13-protofilament (13-pf, blue) microtubules. Top-down views (90° tilt) show the clear arrangement of protofilaments. A region of the 13-pf map is shown with the atomic model of the α/β-tubulin dimer (PDB: 1TUB) docked in**. f.** Distribution of microtubules with 13- or 12-pf across A549-US3 TNTs (n = 17 TNTs, 294 microtubules). **g.** Number of 13- (blue hues) or 12-pf (orange hues) microtubules in ‘plus’ (light) or ‘minus’ (darker) orientation. Each line is an individual TNT. The asterisk (*) denotes data derived from the TNT depicted in (b). **h.** Stacked histogram showing percent microtubules with ‘plus’ orientation ± SD. Circle plot points indicate the percent ‘plus’ microtubules for each individual TNT (n = 17). **i.** Stacked histogram showing percent GFP-EB1 comets with ‘plus’ orientation ± SD. Circle plot points indicate the percent ‘plus’ oriented GFP-EB1 Scale bars: (a) 40 μm; (b) 1 μm; (c) 100 nm.

**Figure 3.**
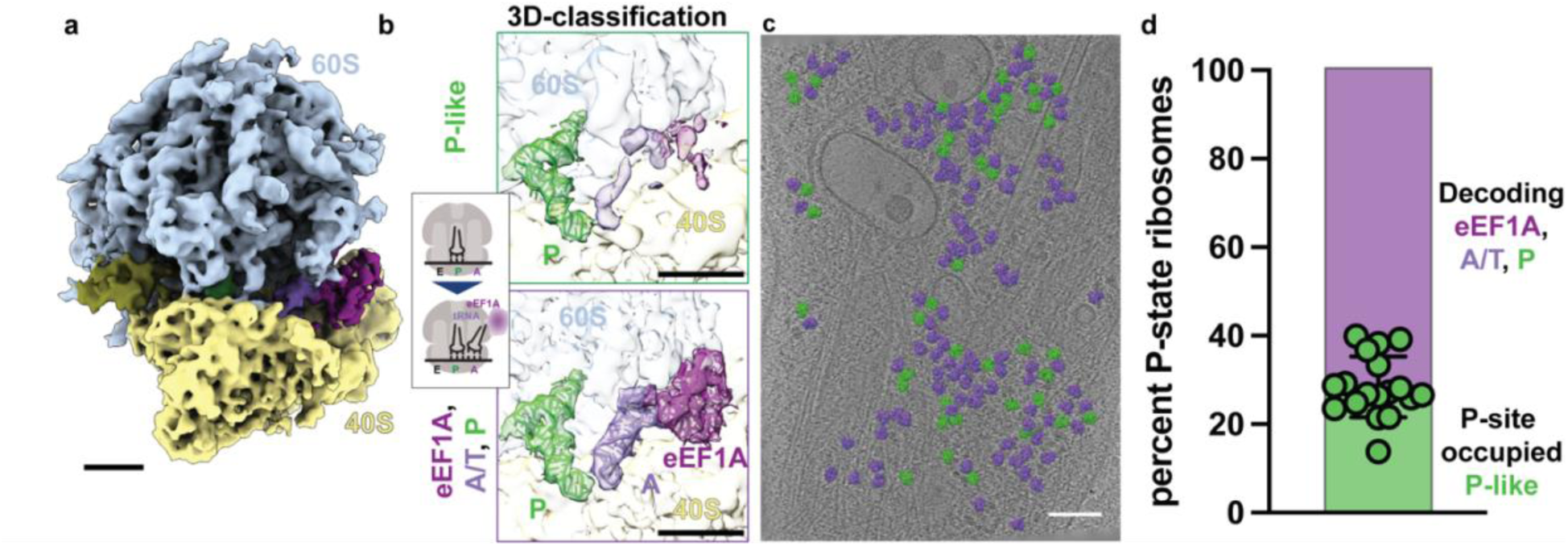
In-situ structural characterization of ribosome functional states within TNTs. **a.** Cryo-ET subtomogram 3-D reconstruction of human ribosomes within the TNT lumen. 60S subunit (blue); 40S subunit (yellow). **b**. Zoom into the tRNA binding sites of maps from two functional states of the elongation cycle (the decoding eEF1A, A/T, P state and the P-state) obtained by 3-D classification. Surfaces correspond to the cryo-ET maps with molecular models docked into them (PDB: 5LZS). Schematic of the corresponding ribosomal elongation cycle states are attached to the zoom-in maps. **c**. Ribosomes in a representative tomogram of a TNT are color-coded by their functional state (Green: P-state; Purple: eEF1A, A/T, P state). **d**. Stacked histogram showing percent ribosomes found in P-state across A549 TNTs (n = 17, 2,402 total ribosomes) ± SD. Circle plot points indicate the percent P-state ribosomes found in each individual TNT. Scale bars: (a-b) 50Å, (c) 100 nm.

Our cryo-ET data revealed macromolecular complexes in the crowded TNT lumen (Fig. 2c; Extended Data Fig. 2g; Extended Data Fig. 3; Extended Data Fig. 4b and d). Microtubules, actin and intermediate filaments ran along the longitudinal axis of the TNT, interspersed with ribosomes, endoplasmic reticulum, and vesicular compartments (Fig. 2c; Extended Data Fig. 3; Extended Data Fig. 4b and 4d). Three-dimensional segmentation demonstrated that these components form a dense network that supports organelle transport and potentially local protein synthesis within these narrow intercellular bridges (Fig. 2c).

**Figure 4.**
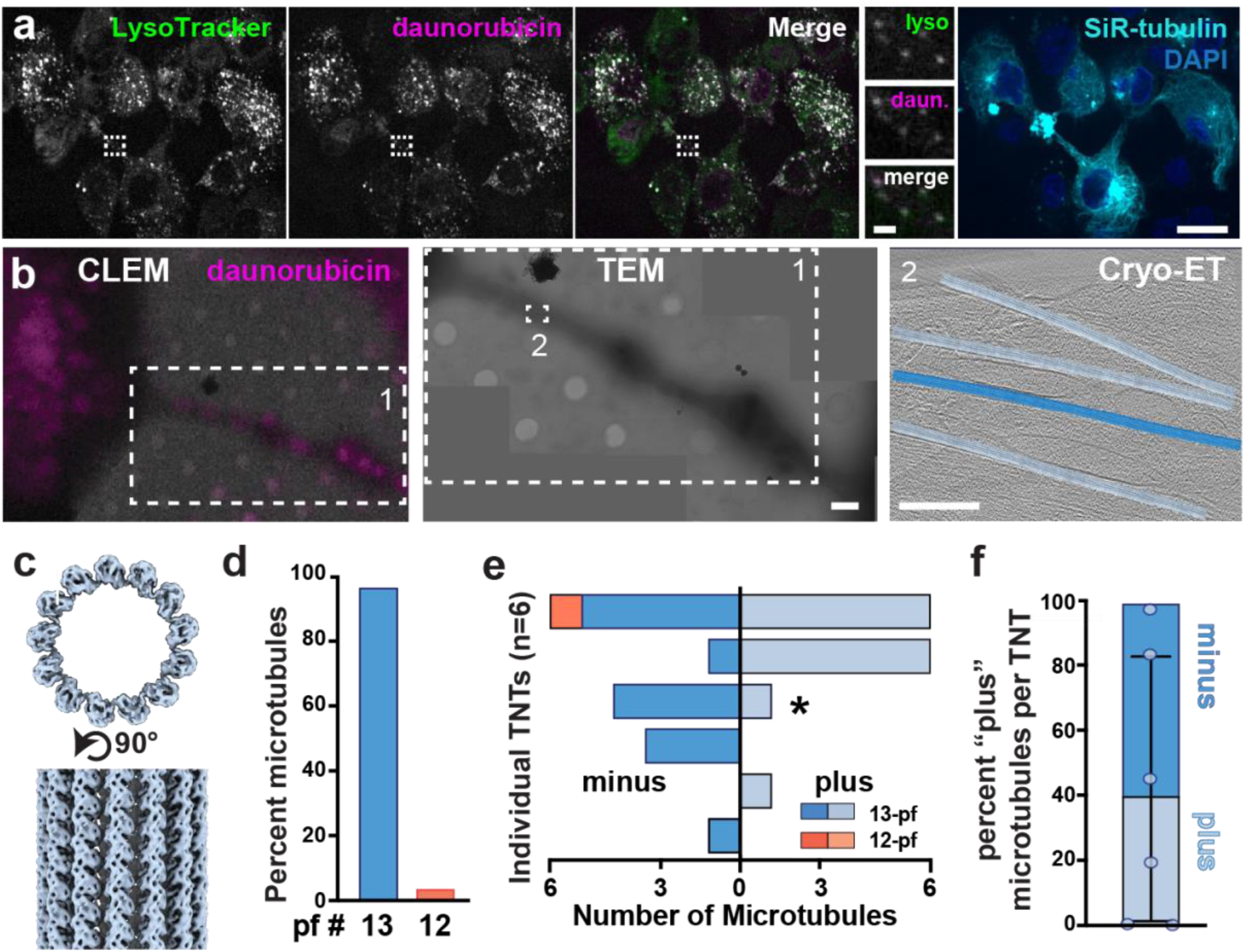
The molecular architecture of TNTs in a chemotherapy-resistance model. **a.** THP-1 cells treated with daunorubicin (magenta), live-stained for lysosomes (LysoTracker; green), tubulin (SiR-tubulin; cyan) and nuclei (DAPI; blue) prior to live-cell imaging (see Supplementary Movie 4). Insets show a magnified view of the boxed area. **b.** CLEM of TNTs in THP-1 cells treated with daunorubicin. (Left) Fluorescence image showing the localization of daunorubicin (magenta) within the lumen of a TNT bridging two cells. (Middle) TEM montage of the targeted TNT. A larger box (1) shows the corresponding areas in the CLEM and TEM views. Smaller white dashed box (2) in TEM image shows a representative tilt-series acquisition area. (Right) tomographic slice (10 nm thickness) of acquisition area (2) outlined in the TEM image. Overlays are microtubule subtomogram averages, reinserted at their original positions and orientations with the same color code as in Figure 2. **c.** 3D subtomogram average of 13-protofilament (13-pf) microtubules from daunorubicin-treated THP-1 TNTs. **d.** Distribution of percent microtubules with 13- or 12-pf across THP-1-daunorubucin treated TNTs (n = 6 TNTs, 29 microtubules). **e.** Number of 13- (blue hues) or 12-pf (orange hues) microtubules in ‘plus’ (light) or ‘minus’ (darker) orientation. Each line is an individual TNT. The asterisk (*) denotes data derived from the TNT depicted in (b). **f.** Stacked histogram shows percent microtubules with ‘plus’ orientation ± SD. Circle plot points indicate the percent ‘plus’ microtubules for each individual TNT (n = 6). Scale bars: (a) 20 μm; (b) 1 μm; (c, f) 100 nm.

### Microtubule architecture and polarity in TNTs

The prevalence of microtubules within TNTs, to our knowledge, provides the first definitive ultra-structural evidence of their presence in these structures. Because microtubules serve as the primary tracks for long-range vesicular transport, we next performed a systematic analysis of their organization. We performed subtomogram analysis of microtubules in TNTs (Fig. 2d). First, we picked subtomograms along each microtubule and obtained subnanometer-resolution maps that aligned with the αβ-tubulin dimer model, enabling confident downstream structural analysis (Fig. 2d-e). We then determined the protofilament number and polarity of individual microtubules. In A549 cells expressing the US3 kinase, we identified a heterogeneous population of microtubule architectures in TNTs. Detailed analysis of 294 individual microtubules revealed that while 13 -pf microtubules are canonical, 12-pf microtubules are a minority population across multiple TNTs (Fig. 2e-f). We reinserted the microtubules with determined polarities back onto the tomograms, which revealed that microtubules within individual TNTs are frequently organized in a bidirectional manner (Fig. 2 e-g). Quantitative analysis across 17 TNTs suggests that TNTs contain microtubules stemming from both cells, providing a structural basis for robust bidirectional cargo trafficking (Fig. 2f and 2g). Because we observed microtubule ends within our tomograms (Extended Data Fig. 4b), we hypothesized that TNT microtubules may be dynamic. Consistent with this, GFP-EB1 comet analysis in US3-induced TNTs confirmed that microtubules within individual TNTs are dynamic (Fig. 2i; Extended Data Fig. 5a-b; Supplementary Movie 3).

### TNT-resident ribosomes are translationally active

The abundance of ribosomes within TNTs suggested that these structures support local protein synthesis (Fig. 2; Extended Data Fig. 2; Extended Data Fig. 4b and 4d). To investigate this, we applied three-dimensional (3-D) classification to resolve the functional states of TNT-resident ribosomes. We identified ∼2,400 ribosomes in TNTs and obtained two major classes: a pre-accommodation decoding state containing eEF1A alongside A/T- and P-site tRNAs, and a “P-like state” where a tRNA occupies the peptidyl (P) site (Fig. 3 a-b) with mostly no occupancy in the A site (POST state). In mammalian cells, the eEF1A-bound decoding state is the most abundant during the elongation cycle, followed by the PRE state (classical A,P), eEF2-bound, and finally the POST state (P-site tRNA only, awaiting next aa-tRNA)^56–58^. The robust presence of the decoding A/T state and a state that is predominantly occupied in the P site (likely composed mostly of POST states but includes decoding, classical PRE and eEF2-bound, combined due to low particle numbers) provides strong evidence that TNT-resident ribosomes are functionally active. Reinserting functional states back into the original tomograms revealed a stochastic spatial distribution (Fig. 3c-d). We did not observe clustering of translation states or tethering to the microtubule cytoskeleton, indicating that active and independent translation occurs along the TNT length.

### TNT molecular landscape in a chemotherapy resistance model

Next, we asked whether TNTs formed under a different biological context are structurally similar. We leveraged the well-established role of TNTs in cancer and modeled TNT function in a chemotherapy resistance model. We used THP-1 cells, a pediatric acute monocytic leukemia cell line, and treated them with daunorubicin, a chemotherapeutic agent commonly used to treat leukemia^59–61^. Daunorubicin contains large aromatic rings and is intrinsically fluorescent, allowing us to track it. We observed daunorubicin inside lysosomes, which were transported bidirectionally through TNTs connecting THP-1 cells (Fig. 4a; Supplementary Movie 4).

Using our CLEM–ET pipeline, we next examined the molecular architecture of these drug-transporting TNTs (Fig. 4b). TEM montages revealed that these TNTs were open (Fig. 4b). Tomography across the TNT length revealed that they exhibited features similar to US3 kinase-induced TNTs (Fig. 4b; Extended Data Fig. 5c-d; Extended Data Fig. 4c and Fig. 2), including TNT thickness (Extended Data Fig. 4a), vesicle content (Extended Data Fig. 4c), and microtubule organization (Fig. 4c-e). Although this dataset was smaller (6 TNTs; 29 microtubules), we obtained a microtubule map with sub-nanometer resolution, enabling detailed downstream analysis (Fig. 4c). Consistent with our observations in US3 kinase-induced TNTs, the microtubules in THP-1 TNTs had predominantly 13 protofilaments (Fig. 4d-e), although 12-pf microtubules were observed in one TNT (Fig. 4e). Microtubule polarity was mixed, mirroring the bidirectional organization seen in US3 kinase-induced TNTs (Fig. 4e-f). These findings indicate that TNTs maintain a conserved molecular and structural landscape across diverse cell types and biological contexts, including under chemotherapeutic stress.

## Discussion

Tunneling nanotubes (TNTs) are thin, long intercellular bridges that mediate the direct transfer of organelles, proteins, and nucleic acids between cells, yet their precise molecular architecture and functional relevance have remained controversial. Prior cryo-ET studies reported actin-based, closed TNT-like structures^3,42^, leaving unresolved whether TNTs are bona fide open conduits supporting bidirectional cargo exchange. While we also observe the same thin actin-rich protrusions, we identify thicker, open, and continuous suspended intercellular bridges. Our comprehensive survey across multiple cell types and biological contexts addresses this critical gap, revealing a rich and conserved molecular architecture comprising actin filaments, microtubules, intermediate filaments, actively translating ribosomes, and diverse organelles including lysosomes, multivesicular bodies, mitochondria, and autophagosomes. We provide definitive evidence that TNTs are continuous open-ended structures capable of sustaining long-range, bidirectional transport. Microtubules within TNTs exhibit mixed polarity, suggesting that both connected cells actively contribute to TNT formation and cargo trafficking. This structural arrangement aligns with observed active lysosomal and drug transport and provides a mechanistic basis for bidirectional cargo movement. Our data also show that TNTs across distinct contexts, namely US3 kinase-induced TNTs in epithelial cells and daunorubicin-transporting TNTs in THP-1 leukemia cells, share remarkably similar molecular architectures, suggesting that TNT formation is a conserved cellular program rather than a context-specific response.

We hypothesize that TNT biogenesis is a cellular stress response: stressed cells emit signals that initiate filopodia-like actin-rich protrusions^3,62^, which upon contact with neighboring cells fuse to form open conduits. Microtubules subsequently invade from both cells, establishing robust bidirectional highways for organelle and vesicle transport. Interestingly, TNT-resident ribosomes appear translationally active, as evidenced by the prominence of ribosomes in a POST state, likely containing only a P tRNA. The enrichment of this POST state in our data suggests that local translation within TNTs may be regulated at the level of tRNA–eEF1A ternary complex availability. Such localized and regulated protein synthesis could support the structural maintenance and functional specialization of these intercellular bridges (Fig. 5).

**Figure 5.**
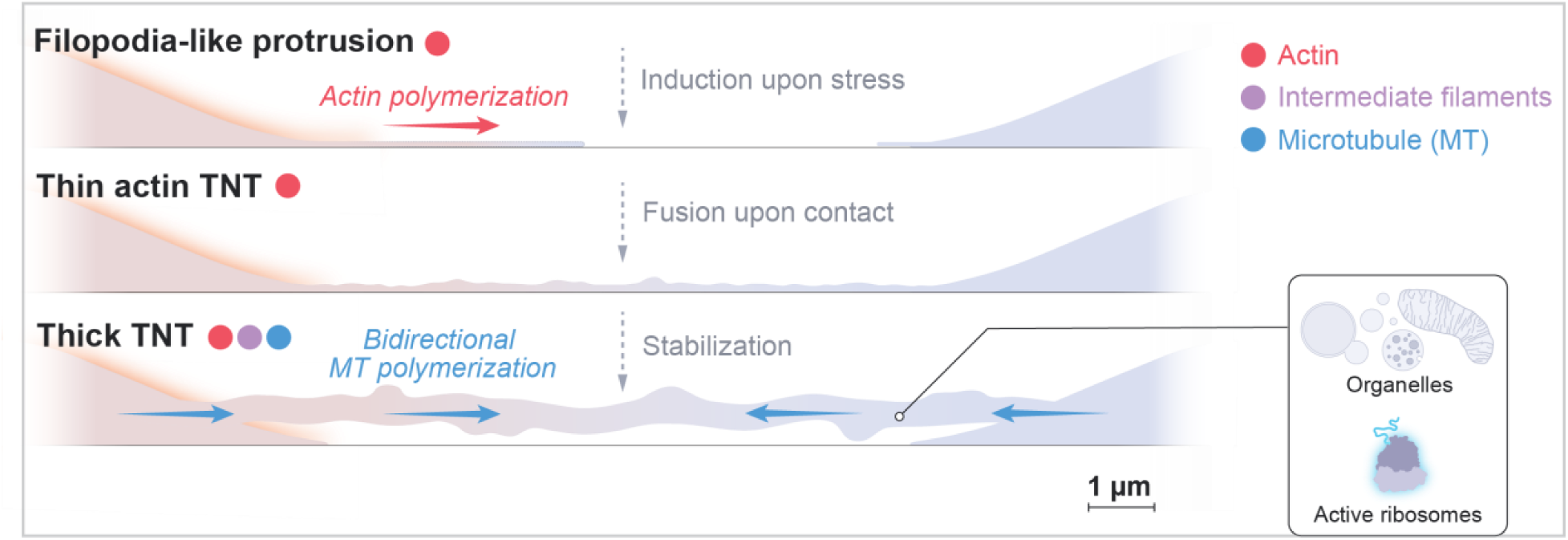
Model of TNT formation, maturation, and cytoskeletal organization. Upon stress, a cell extends an elongated, actin-rich protrusion toward a neighboring cell. Upon contact, this structure fuses with the target cell, establishing a continuous lumen between the two cells. The connection is stabilized as microtubules from both cells invade the TNT bridge, forming an open conduit of bidirectional communication.

This unified model explains TNTs’ broad relevance in diverse physiological and pathological contexts, including embryogenesis, immune cell coordination, neuroprotection, cancer biology, and pathogen dissemination. The ability of TNTs to directly exchange large cellular materials distinguishes TNTs from other intercellular communication mechanisms such as cytonemes, gap junctions, and exocytosis^63^. Our work opens a range of fundamental questions, including how TNTs are initiated, which molecular players drive their formation, how their assembly is coordinated in space and time, and the identities of the nascent proteins translated by TNT-resident ribosomes. Recent studies implicate diverse proteins in TNT initiation ^64–67^, however, a clear consensus is lacking, and how these distinct pathways converge to generate TNTs remains to be determined. Our findings also raise questions about the regulation of transport through TNTs. Despite the structural continuity we document, certain organelles are more abundant than others in TNTs (Extended Data Figures 2-4), but any mechanisms that confer transport selectivity within TNTs remain unknown. Likewise, the molecular machinery governing TNT fusion remains unknown.

While our data support a model in which TNT biogenesis is a conserved, stress-induced program, they also raise a fundamental question: what keeps this program in check? TNT formation in healthy, unstressed tissues must be tightly regulated, as uncontrolled intercellular exchange could have pathological consequences, enabling the spread of toxic cargo, pathogens, or oncogenic material between otherwise insulated cells. The molecular mechanisms that suppress TNT initiation, limit TNT lifetime, or restrict their formation to appropriate cellular contexts remain unknown. Equally unclear is how TNTs are ultimately disassembled. We observed no distinctive structural features that predict which neighboring cells form TNTs and which are bypassed, yet such selectivity likely exists. Identifying the upstream signals that license TNT formation and those that silence it will be a critical next step.

These questions have direct therapeutic relevance. Pharmacological suppression of TNTs could limit tumor microenvironment crosstalk and drug resistance, and block the TNT-mediated dissemination of pathogens, including HIV and SARS-CoV-2. Conversely, enhancing TNT formation or cargo loading could open new avenues for neuroprotection, where TNT-mediated organelle transfer shows promise, as well as for cell-based gene therapy ^27,29^. Collectively, our work provides a molecular-structure framework for TNTs, reconciling previous controversies and laying the foundation for mechanistic and therapeutic exploration of this conserved cellular communication pathway.

## Materials and Methods

### Plasmid construction and lentiviral production

BFP empty vector was purchased from Addgene (plasmid #201383). BFP-US3 and US3-BFP plasmids were synthesized by GenScript, using the BFP plasmid as a backbone. The HTMU lentivirus construct encodes H2B-mCherry-tandem P2A/T2a - US3, cloned in a pFUGW vector. pLenti-HTMU expresses H2B-mcherry (labeling the nuclei of transduced cells) and the *Pseudorabies virus* US3 kinase (untagged) and the construct was generated as follows: US3 from *Pseudorabies virus* strain NIA-3 was ordered as a gBlock with suitable overlapping ends for a 1-fragment Gibson Assembly into the pFUGW-EGFP plasmid. pFUGW-US3 was then digested with BamHI and AgeI and used as the vector to combine a tandem P2A-T2A cleavage site, H2B and mCherry through a 3-fragment Gibson Assembly. Our pLenti-HTMU construct was combined with three packaging vectors (pVSVG, pRSV, pMDL) and were transiently transfected into two 10 cm plates with HEK293T cells. After 48 hours the supernatant of these cultures containing the virus was collected and concentrated with a LentiX Concentrator (Takara Bio). Lentivirus titer was calculated using a LentiX^TM^ GoStix^TM^ (Takara Bio). For transduction, 2.0 × 10⁵ A549 cells were seeded on a 6 well (35 mm). After 24 hours, cells were transduced with a multiplicity of infection (MOI) of ∼0.5.

CTNS-GFP lentivirus was constructed in the self-inactivating lentiviral (SIN-LV) vector backbone and pCCL-EFS-X-WPRE (pCCL), was utilized for all gene transfers. This vector contains the intron-less human elongation factor 1α (EFS) promoter to drive the expression of transgenes, including CTNS-GFP. Lentiviral particles were produced via transient tri-transfection of HEK-293T cells using a third-generation packaging system (pMDLg/pRRE, pHCMV-G, and pRSV-Rev). Supernatants containing the viral particles were collected at 24 hours and 48 hours post-transfection and concentrated by ultracentrifugation at 25,000 rpm for 2 hours at 4°C^28,29^.

### Cell culture and transfection

HEK293T (ATCC; CRL-3216), HeLa (ATCC; CCL-2), primary mouse embryonic fibroblasts, U2OS (ATCC; HTB-96), and A549 (ATCC; CCL-185) cells were cultured in Dulbecco’s Modified Eagle Medium (DMEM; high glucose, 4.5 g/L) supplemented with sodium pyruvate (Corning) at 37 °C and 5% CO₂. Cells were transfected with BFP, BFP-US3, or US3-BFP kinase constructs using Lipofectamine 3000 according to the manufacturer’s instructions. Briefly, 1.6 × 10⁵ cells were seeded in 6-well plates (35 mm) precoated with fibronectin and transfected with 300 ng DNA, 3 µL Lipofectamine 3000, and 3 µL P3000 reagent. Cells were imaged 48 h post-transfection to allow for tunneling nanotube (TNT) formation. THP-1 cells (ATCC TIB-202) were cultured in RPMI-1640 medium (ATCC; 30-2001) supplemented with 10% fetal bovine serum (Gibco One Shot, A3160401) and 0.05 mM 2-mercaptoethanol (Gibco, 21985023) at 37 °C and 5% CO₂. THP-1 cells grow in suspension and were differentiated to enable adhesion for cryo-fixation on EM grids or for live-cell imaging on glass surfaces. For cryo-preservation, a 35 mm glass-bottom dish (Mattek; P35G-1.5-20-C) was fitted with a 15 mm PDMS stencil (Aveole 4W001) containing four 4 mm openings to serve as grid holders. Plasma-cleaned grids (see below) were placed into the stencil wells, coated with fibronectin, and seeded with 75,000–100,000 cells. Cells were differentiated by treatment with 25 ng/mL phorbol 12-myristate 13-acetate (PMA; Sigma-Aldrich; P8139). After 72 h, the medium was replaced with fresh medium containing 100 nM daunorubicin (Sigma-Aldrich; 30450-5MG). Following a 6 h incubation, the medium was exchanged for fresh medium prior to plunge-freezing.

Primary fibroblasts were isolated from neonatal mouse skin biopsies of wild-type (WT) C57B6/J mice. Small skin pieces (approx. 0.5 cm^2^) were attached to culture dishes covered in complete DMEM (high glucose, 4.5 g/L) media supplemented with 10% FBS and 2% penicillin/streptomycin for 3-5 days to allow fibroblasts to grow out of the tissue. Once established, the skin pieces were removed, and the fibroblasts were subcultured ^28,29^.

For the generation of stable cell lines, primary fibroblasts were plated onto fibronectin-coated plates, to enhance transduction efficiency, and transduced with the concentrated virus (MOI ∼20). Successful stable transduction and correct lysosomal targeting of fusion proteins were verified in vitro by observing colocalization with LysoTracker ^29,68^.

All cell lines were routinely tested for mycoplasma contamination by PCR (Myco-Sniff Mycoplasma PCR Detection Kit)

## Cell fixation, immunostaining and live staining

The cells were fixed with prewarmed 3% paraformaldehyde in PHEM (60 mM PIPES, 25 mM HEPES, 10 mM EGTA, 4 mM MgSO₄, pH 6.9; Sigma-Aldrich) for 10 minutes at room temperature. The coverslips were then rinsed twice and washed twice with 1× PBS and quenched with 0.4% NH_4_Cl for 10 min. After washing with PBS, the cells were incubated with blocking and permeabilizing buffer (2% BSA, 0.1% Triton X-100 in 1× PBS) for 20 min at RT. A primary mouse anti-alpha tubulin monoclonal antibody (DM1A; Sigma-Aldrich T6199) was diluted 1:500 in antibody dilution buffer (2% BSA in 1× PBS) and incubated with the cells for 3 h at RT. Following primary antibody incubation, the coverslips were washed three times with 2% BSA in 1× PBS and incubated with 1:200 goat anti-mouse Alexa488 secondary antibody (Invitrogen; A28175, diluted in antibody dilution buffer), and phalloidin 647 (Thermo; A22287) 1:600 for 1 h at RT. Cells were then rinsed five times with 1× PBS before mounting on glass slides using FluorSave (EMD Millipore Sigma). The coverslips were left to dry at least 1 h and imaged immediately or stored at 4 °C for later imaging.

For live-cell imaging experiments, cells were stained with SPY555-tubulin 1 μg/ml or or SiR-tubulin 0.2 μg/m, LysoTracker Green 100 ng/mL dilution and SiR-Actin 0.2 μg/ml. Fibroblasts not transfected with US3 kinase were also stained with Alexa Fluor 405 Wheat Germ Agglutinin (WGA) at 1 μg/ml. Cells were incubated at least 30 minutes before adding 30 mM HEPES pH 7.4 and transferring to the microscope for imaging.

### Confocal fixed and live cell imaging

Fixed cells were imaged on a X-Light V3 50 µm pinholes confocal scanhead (CrestOptics) mounted on a Ti2 Nikon microscope with a Plan Apo lambda 40x NA 0.95 air objective and an automated stage. The microscope was run with NIS Elements using the 488 nm 515 nm and 561 nm and 640 nm lines of a seven-line solid state laser light engine: 405nm, 446nm, 477nm, 520nm, 546nm, 638nm, 749nm (Lumencor Celesta), and an ORCA-fusion digital CMOS camera (Hamamatsu).

Live cell imaging was performed on a Yokogawa W1 confocal scanhead mounted to a Nikon Ti2 microscope with a Plan Apo λ 40X, N.A.0.95 air, a Plan Apo λ 60x NA-1.4 oil and Plan Apo λ 100x NA-1.45 oil objectives (Nikon, Plano Apo). The microscope was run with NIS Elements using the 488 nm 515 nm and 561 nm and 640 nm lines of LUN-F laser engine and two Kinetixs 22 cameras (Photometrics). The scope was equipped with an enclosed temperature incubator to maintain 37 °C. Cells were treated with 30 mM HEPES pH 7.4 prior to imaging to maintain PH and compensate for the lack of CO_2_.

### Cryo–EM sample preparation

Gold Quantifoil R2/2 200–mesh grids or R1/4 200-mesh (Electron Microscopy Sciences) were used for cryo–ET. Prior to sample application, grids were plasma–cleaned for 60 s in a Pelco plasma cleaner using air at 25 W. Immediately after cleaning, grids were coated with 1 μg/mL fibronectin diluted in 1 x PBS and incubated for at least 30 minutes at 37 °C with 5% CO_2_. Cells were then seeded onto grids and left to grow 48 hours for TNT formation. Prior to plunging 4 μl of media was applied to each grid and blotted from the back for ∼6 s using a custom manual plunger. Grids were then plunge–frozen in a 50:50 liquid ethane–propane mixture and stored in liquid nitrogen.

### Cryo-Correlative Light Electron Microscopy (CLEM)

Cryo-Fluorescence microscopy of cells expressing US3 kinase constructs, was performed using an integrated fluorescence microscope (iFLM) system mounted on an Aquilos 2+ (Thermo Fisher Scientific), equipped with a 20 x NA 0.7 air objective and a CoolLED light source for imaging in 4 channels (365 nm/450 nm/550 nm/635 nm). Illumination power was set between 5 % and 10 % with 100 or 150 ms exposure time to minimize photodamage and ∼10 μm Z stacks were acquired at 800 nm steps.

THP-1 cells treated with daunorubicin were imaged with a Leica SP8 point-scanning confocal cryo-CLEM system System equipped with a cryo-stage maintained by nitrogen air flow, enabling high-resolution cryo-correlative imaging. The microscope is an upright, fixed stage system equipped with a 20X 0.75 NA air objective, four laser lines at 405 nm/488 nm/561 nm /633 nm, and 4 detectors (two GAsP/APD HyD high sensitivity detectors and two photomultiplier tube (PMT) detectors). ∼10 μm Z stacks were acquired with a 500 nm step size with 10 % laser power and 200 ms exposure.

### Cryo–ET data acquisition

For all datasets collected on TNT, a Titan Krios G3 transmission electron microscope (Thermo Fisher Scientific) operating at 300 kV and equipped with a K3 direct electron detector (Gatan) was used for data acquisition. A nominal magnification of 53,000× (1.635 Å pixel size, counting mode) was used during tilt–series collection, with a 100 μm objective aperture inserted. A zero–loss energy filter with a 15 eV slit width was applied throughout imaging.

For sample navigation, overview images were first acquired on TNT at a nominal magnification of 8,700× (pixel size 10.64 Å, counting mode) at 0 ° and montaged for tomogram targeting. Targets were selected manually while avoiding regions that were too dark for reliable imaging.

Tilt–series data were collected automatically using paceTOMO ^69^ implemented in SerialEM ^70^ (http://bio3d.colorado.edu/SerialEM/, RRID:SCR_017293). A dose-symmetric tilt scheme from – 37.5° to +37.5° with 2.5° increments was used, yielding 31 tilts per series. Each tilt consisted of 8 frames. The total electron dose was 160 e−/Å² with a dose rate of ∼35 e−/Å²/s, and the defocus was set to –3 to –5 μm. Data collection parameters are summarized in Supplementary Table 1.

### Tilt–series preprocessing and tomogram reconstruction

Warp^71^ (http://www.warpem.com/warp/, RRID:SCR_018071) was used for initial pre–processing of the tilt series, including motion correction, CTF estimation, and gain correction. Tilt–series alignment was then carried out in eTomo within the IMOD package^70^ (v4.11.25, https://bio3d.colorado.edu/imod/, RRID:SCR_003297) using patch tracking in an 11×8 grid. Residual tracking errors were calculated by eTomo, and a custom Python script was used to remove patches or tilts whose errors exceeded two standard deviations within their respective distributions, thereby improving the mean residual alignment error. The refined alignment parameters were exported as xf files and imported back into Warp for tomogram reconstruction using weighted back–projection, with tomograms binned to 10 Å per pixel. Deconvolution was performed in Warp using default settings (high–pass 300, strength 1, fall–off 1). For visualization in the figures, tomograms were either reconstructed in IMOD using SIRT (Simultaneous Iterative Reconstruction Technique) for 10 iterations, or optimized alignment using miss-alignment^72^, then denoised using IsoNet2^73^. A Python–based wrapper package that automates and streamlines the entire processing workflow has been made available at https://github.com/Siyu-C-TOMO/Warp_Pipeline.

### Microtubule tracing and protofilament number determination

IMOD was used to visualize deconvolved tomograms and manually trace microtubule backbones. Using the low–magnification TNT overview as a reference, microtubules were consistently picked from right to left to ensure that the skewing pattern observed in class averages could be unambiguously related to microtubule growth polarity. The coordinates of the traced microtubules were imported into Dynamo (running in MATLAB, v11555, https://www.dynamo-em.org/w/index.php?title=Main_Page, RRID:SCR_025592)^74^ where they were resampled at equal spacing for subtomogram extraction. To minimize missing–wedge bias, the azimuthal angle around the microtubule axis was randomized for each subtomogram, as performed previously^55,75,76^.

Subtomograms were first aligned to a featureless cylindrical reference, using local angular and translational constraints limited to the first two Euler angles and movements along the microtubule axis for a single iteration (box size 40 nm). To determine the protofilament number, aligned subtomograms from each microtubule were subjected to five iterative alignments with enforced symmetries ranging from C11 to C16. Resulting averages were maximum–intensity projected along the Z–axis, and the symmetry producing the sharpest protofilament features was selected to determine protofilament number. Because microtubules in tomograms adopt arbitrary orientations, subtomograms displaying opposite skewing directions were identified and flipped in Dynamo to maintain consistent polarity. Duplicate subtomograms were then removed based on distance. Microtubules that did not yield an unambiguous protofilament number were excluded from further analysis.

Subtomograms corresponding to 12-pf and 13-pf microtubules were then re–extracted separately and subjected, in parallel to three iterations of local refinement in Dynamo without imposing symmetry, yielding improved microtubule averages. The resulting coordinates and particle stacks were subsequently used for downstream subtomogram analyses in Relion3 ^77^ (http://www2.mrc-lmb.cam.ac.uk/relion, RRID:SCR_016274) and in M (integrated within Warp)^71^.

### Microtubule subtomogram analysis

For 3D reconstruction of microtubules from A549–US3 TNTs, 29,514 subtomograms corresponding to 294 microtubules across 17 TNTs were extracted in Warp and imported into Relion3. Subtomograms were at bin-4 (6.54 Å per pixel; box size 72), and metadata included microtubule identifiers (HelicalTubeID) and orientation information, with the AnglePsiFlipRatio parameter set to 0.9 to suppress unintended polarity flipping. A single–iteration 3D classification using initial references representing 11-pf to 16-pf architectures^78^ assigned 81% of particles to the pf–13 class, supporting the accuracy of the protofilament number determined in Dynamo (Extended Data Fig. 6). Refinement was then performed with C1 symmetry and a cylindrical mask, followed by local alignment using helical parameters (angular search range 21.6° with 1.8° steps, twist 27.7° to correct for mirror–handedness, and rise equivalent to 9.36 Å in binned subtomograms).

To improve map quality, aligned subtomograms were subjected to six-class 3D classification without symmetry or alignment. To prevent overfitting, the regularization parameter T was set to 1 instead of default 4, and the resolution during the E–step was limited to 15 Å, effectively disabling high–frequency alignment. The sub-class exhibiting the clearest structural features (10,653 subtomograms) was re–extracted at 3.27 Å per pixel (box size 144) and refined to a final resolution of 12.4 Å. Using an analogous workflow, 3,183 subtomograms from 12-pf microtubules (32 microtubules) were refined to 18 Å without 3D classification, using a twist of 29.88° and rise equivalent to 10.38 Å in binned subtomograms.

Refined subtomogram coordinates from both protofilament populations were then processed as separate groups within the same refinement pipeline in Warp/M. Iterative subtomogram refinement—comprising particle alignment, image–warp correction, defocus optimization, and refinement of higher–order CTF and geometric parameters, yielded final resolutions of 8.0 Å for 13-pf microtubules and 13.3 Å for 12-pf microtubules. For each subtomogram, only the central slice along the MT axis (30% of the box, parameter Central Z length) was used for alignment. All reported resolutions correspond to the gold–standard Fourier Shell Correlation at the 0.143 criterion^79,80^.

A similar workflow was applied to the THP–1 TNT dataset, comprising 2,920 subtomograms from 27 13-pf microtubules. As only two 12-pf microtubules were identified, 12-pf refinement was not pursued. Warp/M refinement produced an 8.7 Å reconstruction of 13-pf microtubules (Extended Data Fig. 7; Supplementary Table 1) . UCSF ChimeraX (v1.10.1, https://www.cgl.ucsf.edu/chimerax/, RRID:SCR_015872) was used for volume segmentation, figure and movie generation, and rigid-body docking^81^.

### Ribosome subtomogram analysis

Ribosomes from A549–US3 TNTs were first identified by 3D template matching in Warp. Z–boundaries of each tomogram were automatically detected using IMOD’s *findsection* tool and manually refined to remove off–targets. A manually chosen cross–correlation threshold was applied to exclude low–confidence matches, and the remaining coordinates and initial orientations were imported into Relion3 for subtomogram extraction (bin–4; pixel size 6.54 Å; box size 72). A six–class 3D classification using default alignment parameters (100 iterations; regularization parameter T=1; angular step 7.5°; shift range 5 pixels, step 1 pixel) was performed to remove false positives. The resulting 1,803 subtomograms were refined, re–extracted at bin–2 (box size 144), and locally refined to 12.4 Å. Subsequent M–refinement yielded an overall resolution of 9.1 Å. To improve picking completeness, the refined coordinates were converted into crYOLO^82^–training CBOX files using our custom Python workflow, in which Z–positions were expanded at 10–pixel intervals to ensure full volumetric sampling. After applying an identical cleaning procedure and removing duplicates from the combined particle set, an additional 599 subtomograms were retained for a total of 2,402 ribosomes for classification. To classify ribosome translation states, a six–class 3D classification without alignment separated two ribosomal functional states, yielding 1,715 decoding-state and 687 P-like state particles. This latter map has a clear P-state density, but still contains density in the A site at low threshold, indicating that it may contain A/A and A/T tRNAs and eEF2. These two particle groups were refined independently within the same Warp/M pipeline, producing reconstructions at 10.2 Å (decoding state) and 13.1 Å (P-like state) (Extended Data Fig. 8; Supplementary Table 1). All custom code is available through the Warp_Pipeline repository referenced above and the accompanying star–handler package (https://github.com/Siyu-C-TOMO/star_handler).

### Model building

Rigid–body fitting was used to generate all microtubule and ribosome models. The α/β–tubulin dimer (PDB 1TUB) and the human ribosomal elongation complex (PDB 5LZS) were docked into the corresponding subtomogram–averaged maps using UCSF ChimeraX without additional manual rebuilding or flexible refinement (Supplementary Table 1).

### Tomogram Annotation and 3D Modeling

Microtubule backbones were traced in IMOD, and their polarity and orientations obtained from subtomogram analysis were applied during tomogram–level annotation. Subtomogram lists, particle coordinates, orientations, and class averages of 13-pf and 12-pf microtubules, as well as ribosomes, were imported into ArtiaX^83^ in UCSF ChimeraX for spatial placement and visualization within each tomogram. Vesicular membranes were segmented using MemBrain–seg^84,85^ with the best–trained model and default settings, followed by manual refinement and color–coding in ChimeraX. All annotated structures were assembled into a unified coordinate frame to produce 3D visualizations of the spatial organization in individual tomograms.

For geometric analysis, microtubules in each TNT were grouped by protofilament architecture (12-pf or 13-pf) and by orientation (plus or minus). These values were used to compute microtubule polarity histograms and protofilament–occupancy ratios.

### Data and statistical analysis

Fluorescent images were processed in Fiji^86^. To quantify the percentage of cells forming TNTs, we identified connecting actin and membrane lines between cells. Very thin processes usually < 4 μm long were not included. Mitotic cells (including cytokinetic cells) were also excluded from our analysis; cytokinetic intercellular bridges are readily distinguishable due to their distinct tubulin stain. Each TNT was tracked with the line tool to measure TNT number and length. Circle ROIs were used to annotate cells. In Excel, the number of cells, TNTs and TNT length was tracked. For each TNT, we also noted whether it was positive for actin only, or actin and tubulin staining.

In our cryo-ET analysis, all TNTs were treated as independent biological units. For ribosomes, the P–state ratio per TNT was calculated as the number of P–state ribosomes divided by the total ribosome number. All quantifications were performed per TNT and reported as aggregate distributions across the dataset.

## Data availability

Subtomogram-averaged maps of the 13-pf microtubule, 12-pf microtubule, and the *in situ* ribosome states generated in this study have been deposited in the Electron Microscopy Data Bank (EMDB) under accession codes EMD-XXXX, EMD-YYYY, and EMD-ZZZZ. Source data for cell biological analyses, tomogram thickness and vesicle diameter are provided with this paper. All other data supporting the findings of this study are available from the corresponding authors upon reasonable request.

## Code availability

The Python-based wrapper package for the automated Warp processing workflow is available at https://github.com/Siyu-C-TOMO/Warp_Pipeline. The accompanying custom star-handler package is available at https://github.com/Siyu-C-TOMO/star_handler.

## Acknowledgements

S.R.P. and E.V. are Investigators of the Howard Hughes Medical Institute. The authors acknowledge the facilities of the UC San Diego cryo-EM facility, part of the Goeddel Family Technology Sandbox, along with the scientific and technical assistance of facility staff members Dr. Mariusz Matyszewski and Dr. Inga Kuschnerus. We also thank Andres Leschziner for helpful discussions that informed our imaging scheme and data processing pipelines. Molecular graphics and analyses were performed in part using UCSF ChimeraX, developed by the Resource for Biocomputing, Visualization, and Informatics at the University of California, San Francisco, with support from NIH R01–GM129325 and the Office of Cyber Infrastructure and Computational Biology, National Institute of Allergy and Infectious Diseases. We also thank the UC San Diego Physics Computing Facility for providing computational support. This work was also supported by the National Science Foundation (DBI-1920374 to E.V.). S.C. and E.K. were supported by the Jane Coffin Childs Memorial Fund for Medical Research. T.B. was supported by an American Heart Association Predoctoral Fellowship. J.H. was supported by an EMBO Long-Term Fellowship (ALTF 871-2020).

## Author contributions

Conceptualization: S.R.P., E.K., E.V.

Methodology, Data Collection, Visualization and Analysis: E.K., Si.C., F.S., R.A.B.-G., R.G.A.,

M.R. A.F., L.V.N., N.L.

Writing: E.K., Si.C wrote the original draft. Review & Editing: E.K., Si.C., E.V., S.R.P., with input from all authors.

---

**Extended Data Fig. 1.**
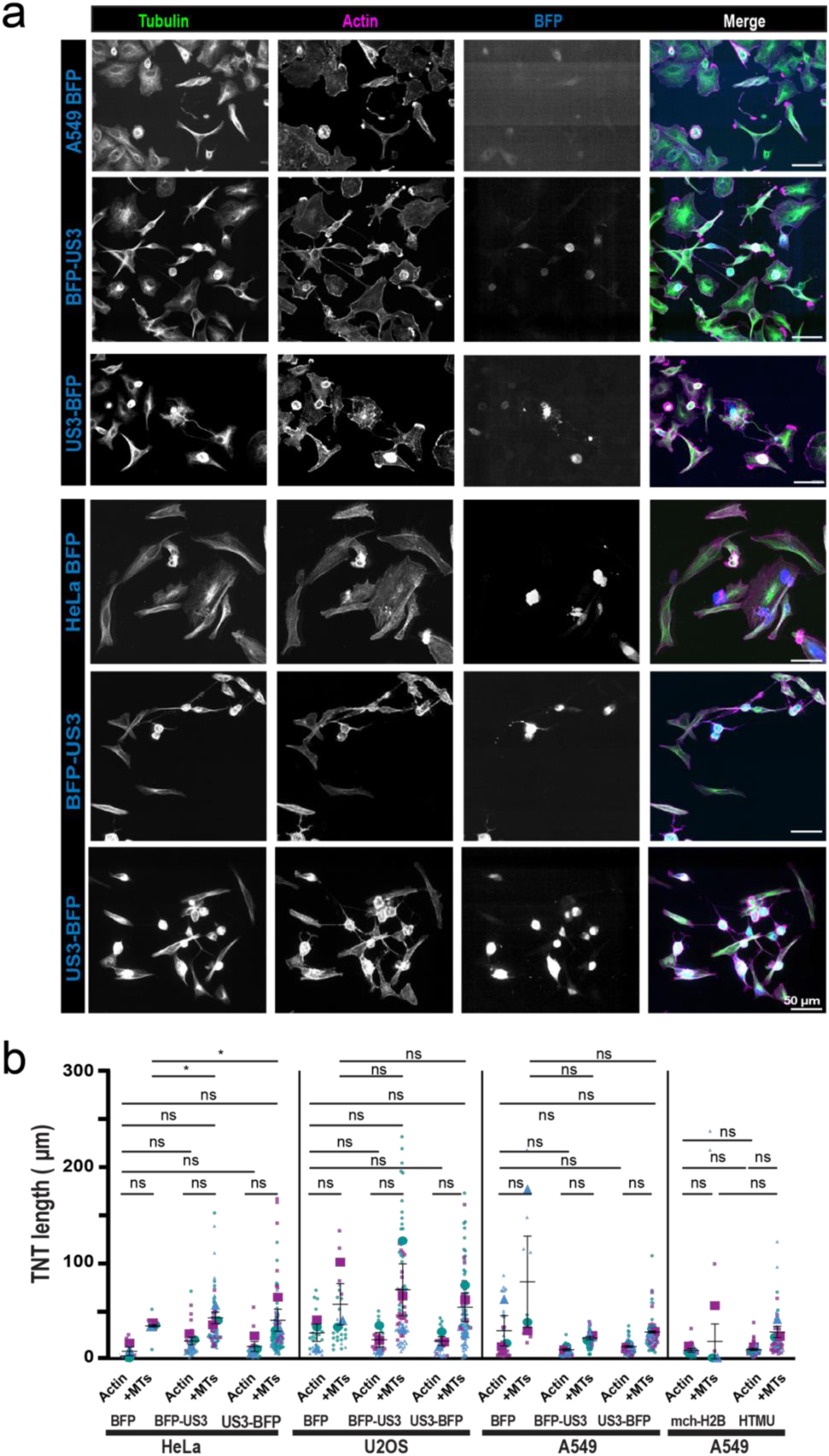
US3 kinase induces TNT formation in different cells lines. **a.** Gallery of representative images of A549 or HeLa cells expressing BFP, BFP-US3 or US3-BFP constructs, and stained for actin (phalloidin) and microtubules (DM1α). **b.** Quantification of TNT lengths that were positive only for an actin stain (phalloidin 647), or for both actin and microtubules (DM1α; Sigma), across different cells lines. Superplots show all individual data points for each of the three biological replicates. Larger opaque shapes show the mean of each biological replicate. Βar graph and error bars show the mean ± SD. HeLa BFP: n= 3/2/1 actin TNTs, n= 0/4/4 +MTs TNTs, HeLa BFP-US3: n=28/22/15 actin TNTs, n= 7/16/22 +MT TNTs, HeLa US3-BFP: n= 62/24/25 actin TNTs, n= 15/7/11 +MT TNTs. U2OS BFP: n= 18/5/8 actin TNTs, n= 18/2/6 +MT TNTs, U2OS BFP-US3: n= 30/31/21 actin TNTs, n= 8/16/7 +MT TNTs, U2OS US3-BFP: n=34/22/38 actin TNTs, n= 9/8/11 +MT TNTs. A549 BFP: n= 2/3/5 actin TNTs, n=5/11/11 +MT TNTs, A549 BFP-US3: n= 31/15/16 actin TNTs, n= 14/18/20 +MT TNTs, A549 US3-BFP: n=26/21/11 actin TNTs, n= 23/27/15 +MT TNTs. A549 H2B-mcherry: n=11/10/6 actin TNTs n= 0/2/0 +MT TNTs, A549 HTMU: n= 12/26/11 actin TNTs, n= 28/41/16 +MT TNTs TNTs quantified from the >200 cells per replicate in Fig.1e. MT; microtubule

**Extended Data Fig. 2.**
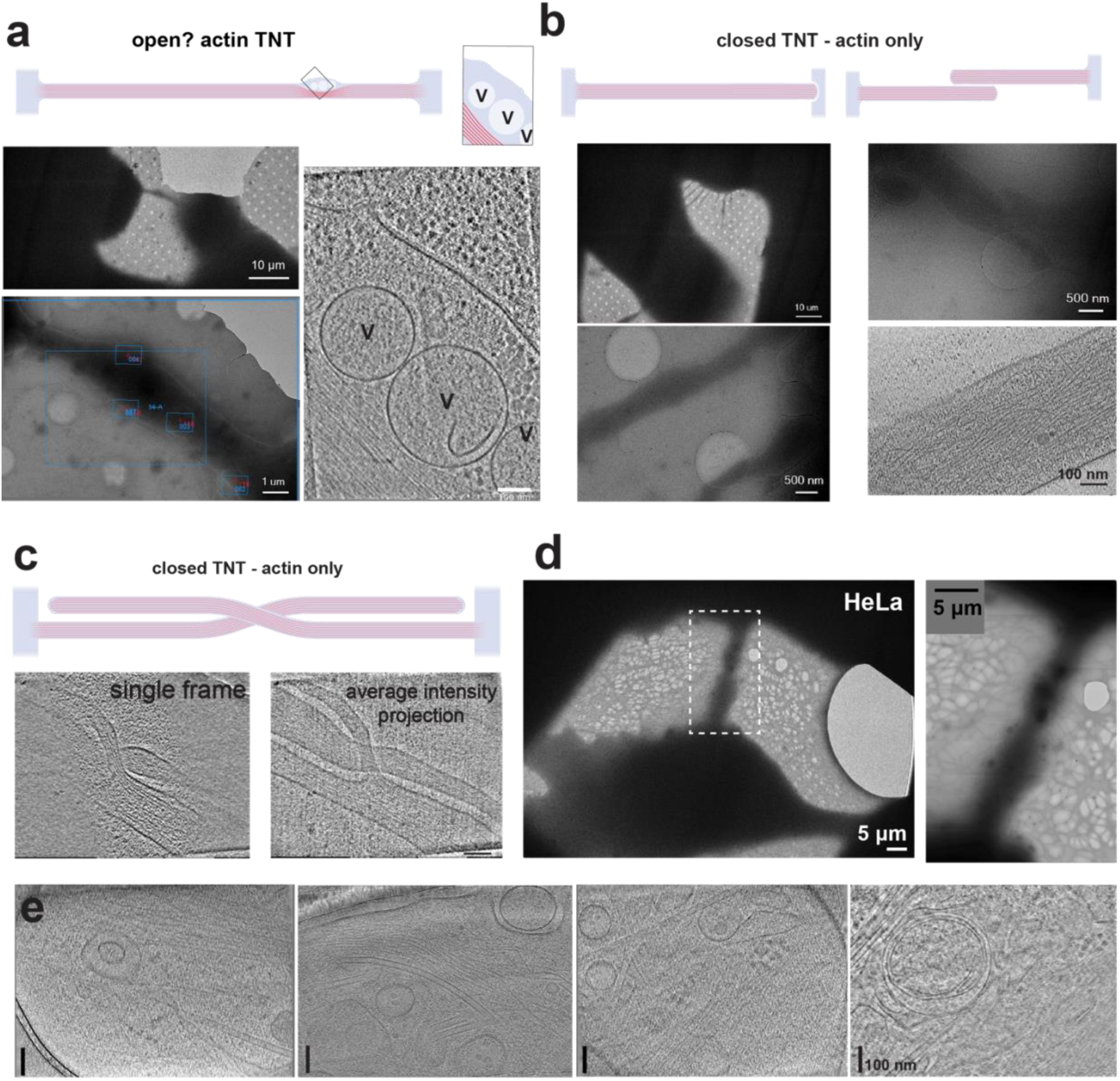
Gallery of TEM images of thin and HeLa TNTs. **a-c**. Representative examples of thin actin-based TNTs shown at low magnification (grid squares and montages), or as higher magnification tomograms. TNTs may appear potentially open (a) or closed-ended (b-c). Vesicular structures are occasionally observed in a subset of thin actin TNTs (a). **d**. TEM images of open TNTs in HeLa cells expressing US3–BFP. **e**. Tomograms acquired from TNT shown in d. Scale bars in tomogram slices (a, b, c and e) are 100 nm.

**Extended Data Fig. 3.**
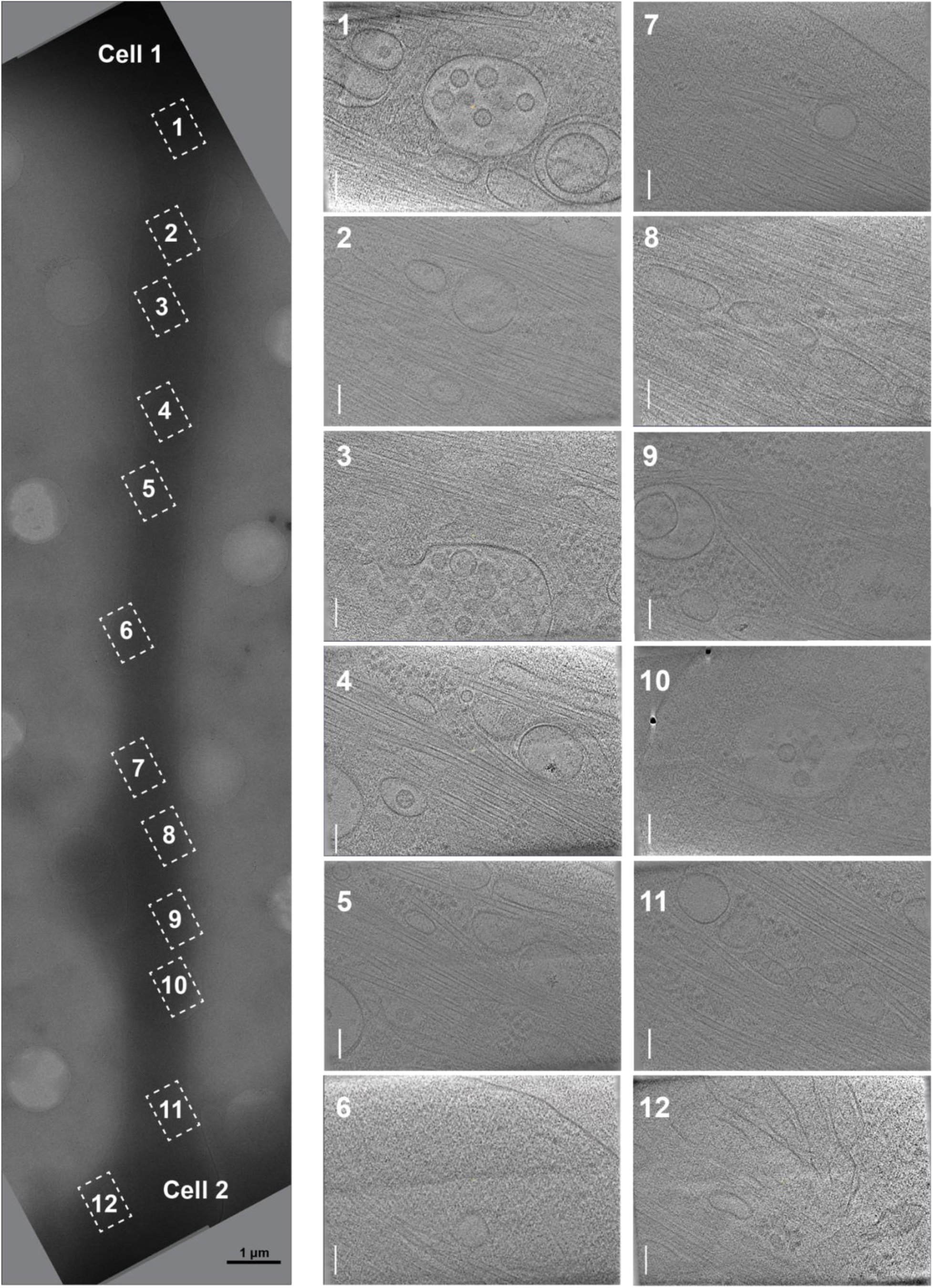
TEM images across an open TNT. Close-up of the open TNT shown in Fig. 2b. Reconstructed tomogram slices (10 nm thickness) are overlaid on all imaged regions. These include the tomograms used for microtubule (MT) reconstruction in Fig. 2a. Scale bars in tomogram slices are 100 nm.

**Extended Data Fig. 4.**
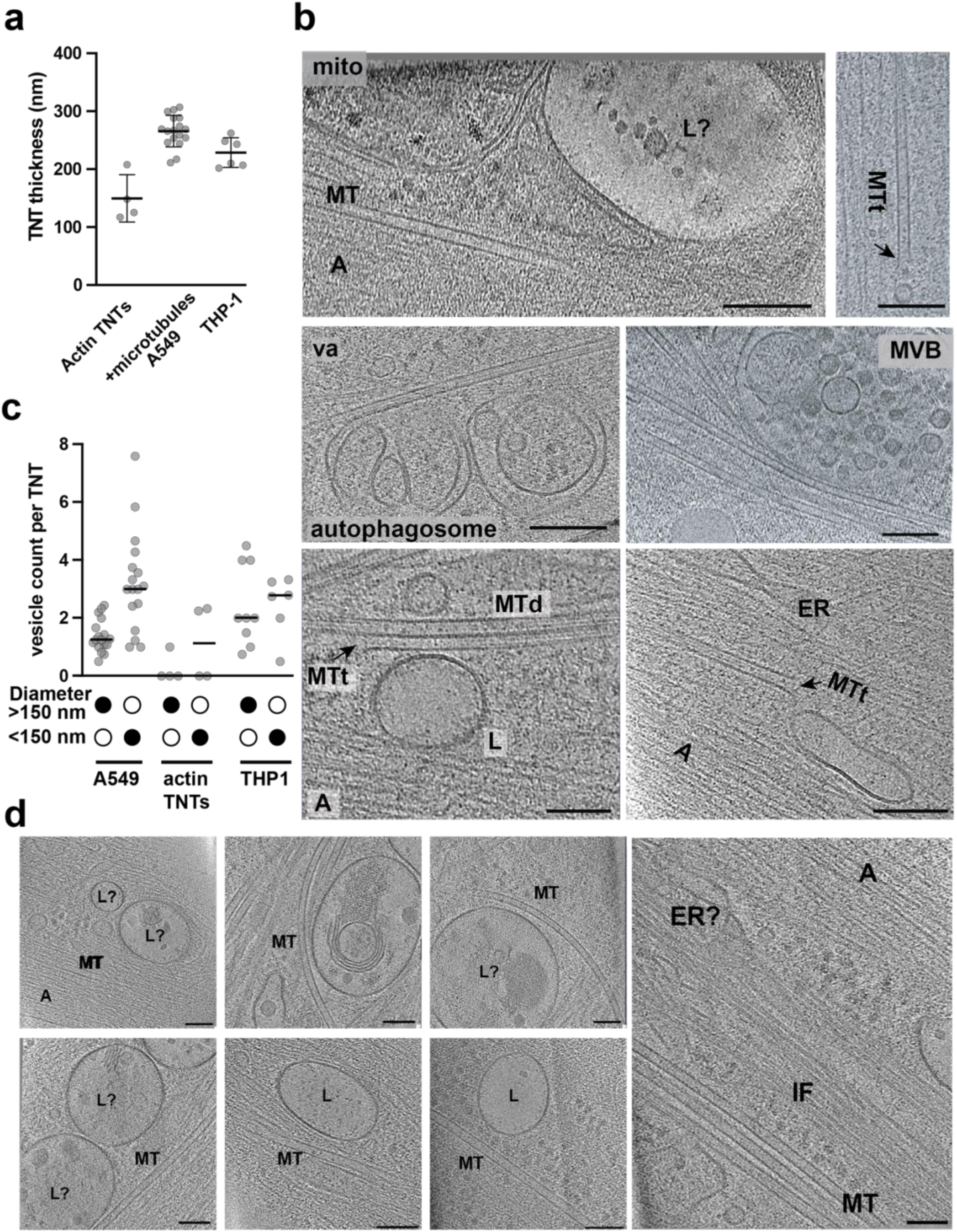
Gallery of tomograms obtained from A549-US3 TNTs. **a.** Quantification (mean ± SD) of tomogram thickness per TNT. Each datapoint represents the average thickness of each TNT, generated by measuring the thickness of all tomograms acquired across the length of the TNT. A549; n=17, THP-1 n=6 TNTs, thin actin TNTs n= 4 **b.** Gallery of tomograms showing the different vesicles, cytoskeleton and cellular materials found in TNTs. Scale bars are 100 nm. **c.** Quantification of number of vesicles with diameter >150 nm and <150 nm per TNT in A549 cells, THP-1 cells or actin only TNTs. A549; n=17, THP-1 n=6 TNTs, actin TNTs n= 4. Bar denotes the mean. **d.** Gallery of additional tomograms found in open TNTs. Scale bar 100 nm. MT; microtubule, MTt; microtubule tip, MTd; microtubule doublet, va; vault, A; actin, IF; intermediate filament, ER; endoplasmic reticulum MVB; multivesicular body, L; lysosome. mito; mitochondrion.

**Extended Data Fig. 5.**
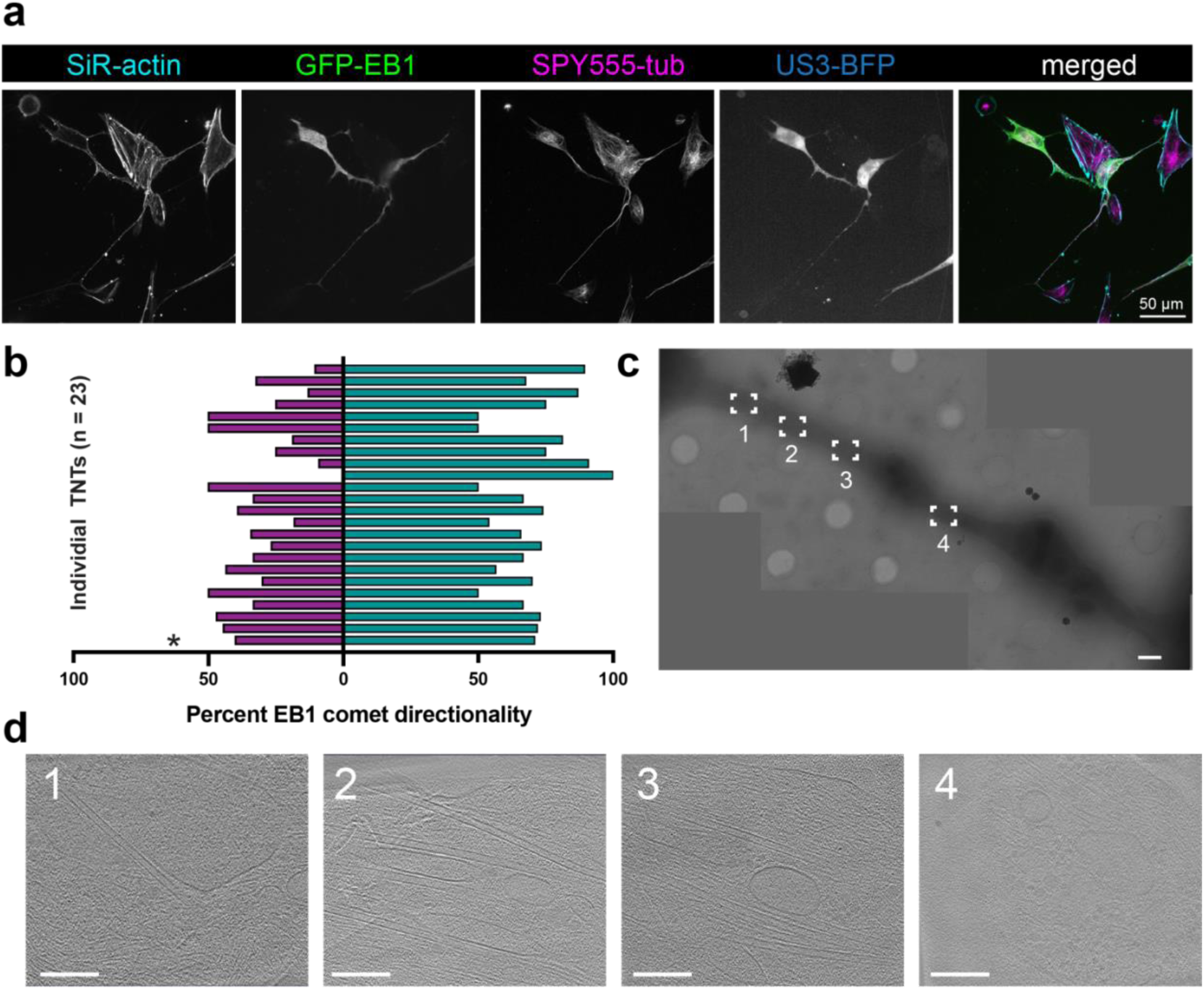
TNTs contain dynamic, mixed polarity microtubules and THP-1TEM data acquisition areas. **a.** Representative image of a tunneling nanotube formed between HeLa cells expressing US3 - BFP and GFP-EB1. Cells were live-stained with SiR-actin and SPY555-tubulin. **b.** Plot showing percent EB1 comets moving in one direction (plum) or the other (teal). Each line is an individual TNT. The asterisk (*) denotes data derived from the TNT depicted in a. **c.** TEM montage of the open TNT in THP-1 cells shown in Fig. 4b. Boxed regions indicate sites of tomogram acquisition. **d.** Tomogram slices (10 nm thickness) acquired from TNT shown in c. Scale bars are 10 μm in (a), 2 μm in (c) and 100 nm in (d).

**Extended Data Fig. 6.**
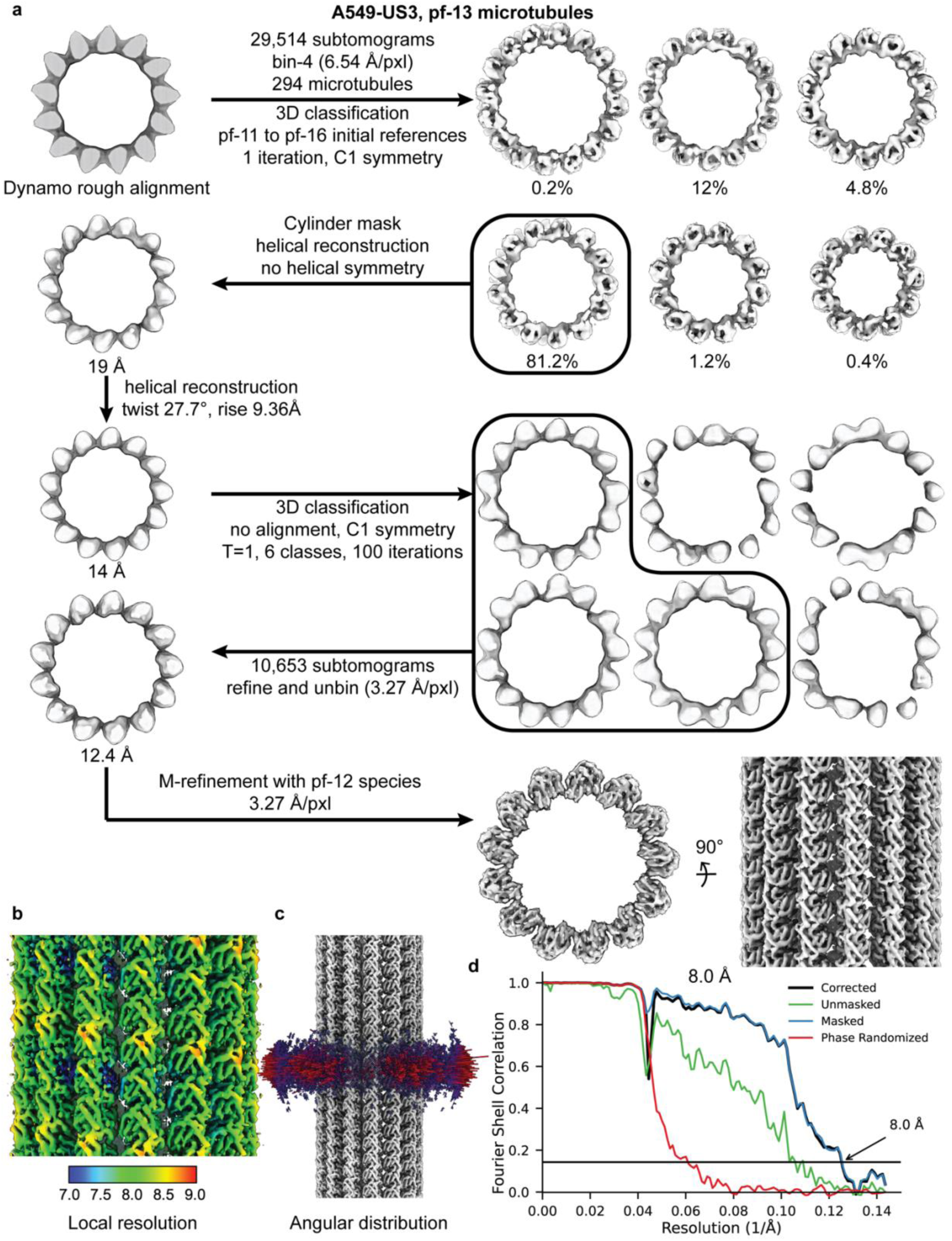
Subtomogram averaging workflow and resolution assessment of A549-US3 13-pf microtubules. **a.** Data-processing workflow for A549-US3 13-pf microtubules, from rough alignment and classification to final refinement. Representative end-on and side views of intermediate and final reconstructions are shown. **b.** Local-resolution map of the final 13-pf microtubule reconstruction. **c.** Angular distribution of subtomograms used for the final reconstruction. **d.** Gold-standard Fourier shell correlation (FSC) curves of the final reconstruction; the masked FSC reached the 0.143 criterion at 8.0 Å.

**Extended Data Fig. 7.**
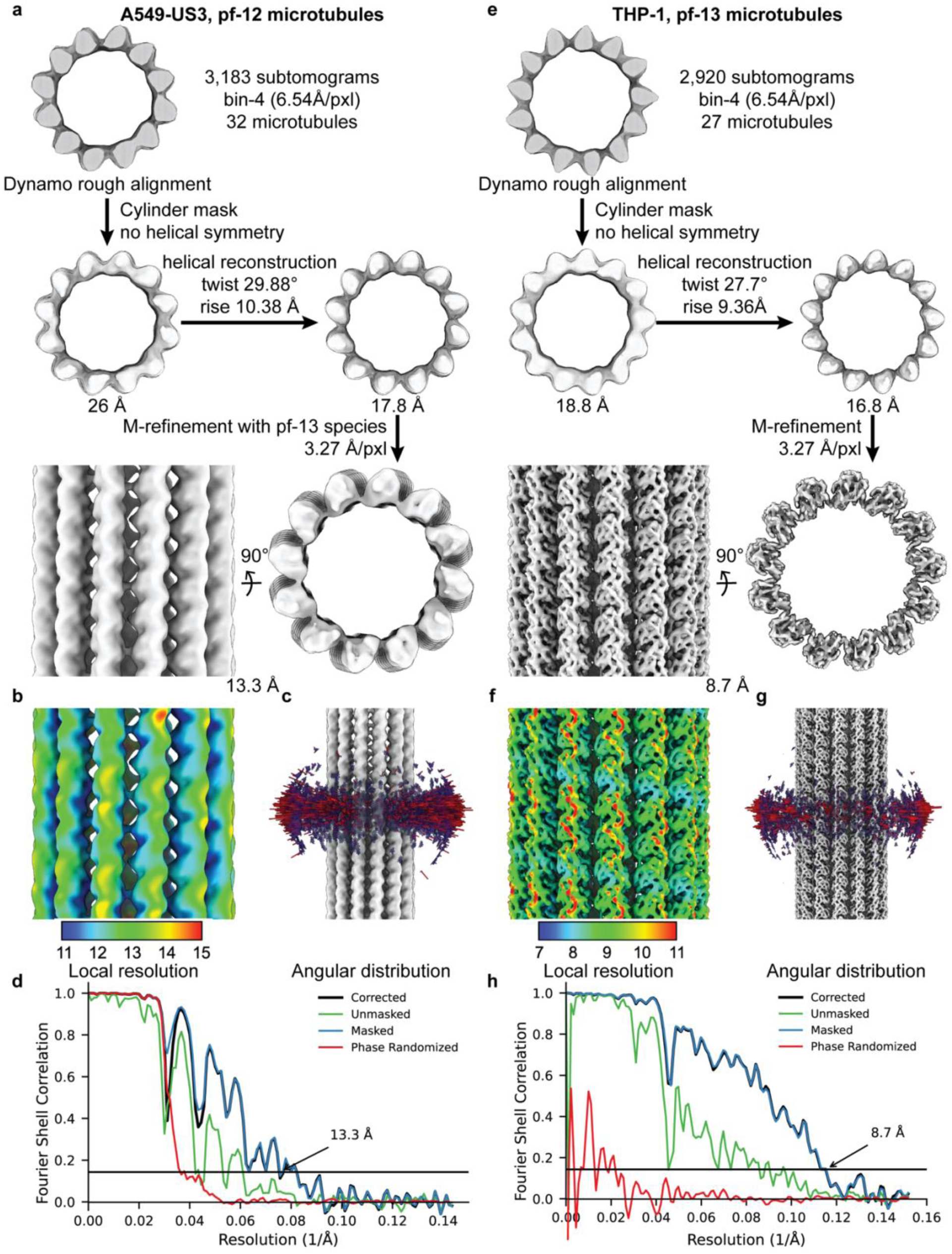
Subtomogram averaging and validation of A549-US3 12-pf and THP-1 13-pf microtubules. **a.** Data-processing workflow for A549-US3 12-pf microtubules, from rough alignment to final M-refinement. Representative side and end-on views of the reconstruction are shown. **b.** Local-resolution map of the final A549-US3 12-pf reconstruction. **c.** Angular distribution of subtomograms used for the final A549-US3 12-pf reconstruction. **d.** Gold-standard FSC curves of the A549-US3 12-pf reconstruction; the masked FSC reached the 0.143 criterion at 13.3 Å. **e.** Data-processing workflow for THP-1 13-pf microtubules, from rough alignment to final M-refinement. Representative side and end-on views of the reconstruction are shown. **f.** Local-resolution map of the final THP-1 13-pf reconstruction. **g.** Angular distribution of subtomograms used for the final THP-1 13-pf reconstruction. **h.** Gold-standard FSC curves of the THP-1 13-pf reconstruction; the masked FSC reached the 0.143 criterion at 8.7 Å.

**Extended Data Fig. 8.**
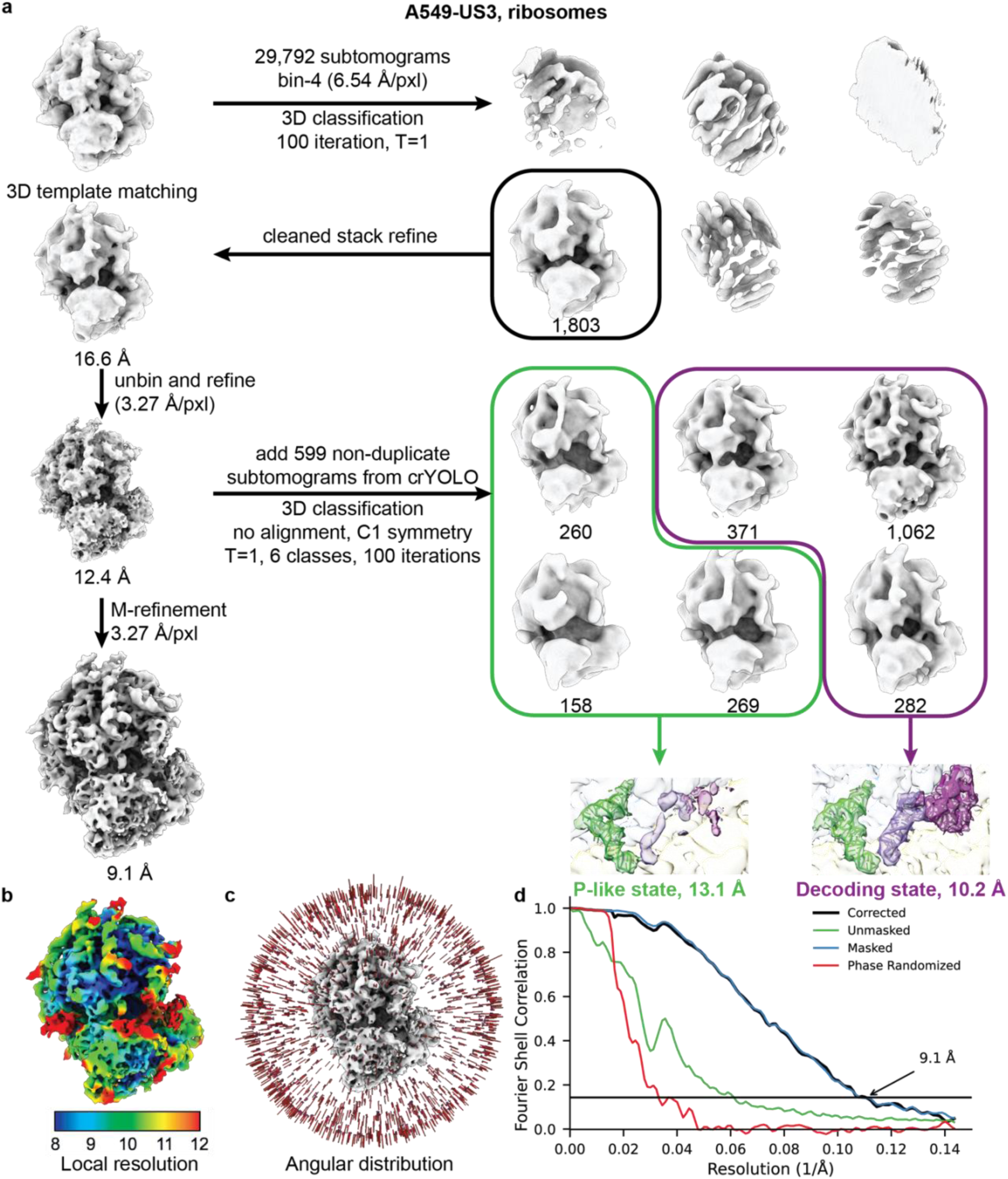
Subtomogram averaging, classification and resolution assessment of A549-US3 ribosomes. **a.** Data-processing workflow for ribosome subtomogram averaging in A549-US3 cells, from 3D template matching and classification to final refinement. Additional non-duplicate particles identified by crYOLO were included for no-alignment 3D classification, yielding ribosome classes corresponding to the P-like state and decoding A/T state. Representative reconstructions of the overall ribosome map and the classified states are shown. **b.** Local-resolution map of the final ribosome reconstruction. **c.** Angular distribution of subtomograms used for the final ribosome reconstruction. **d.** Gold-standard FSC curves of the final ribosome reconstruction; the masked FSC reached the 0.143 criterion at 9.1 Å.

**Supplementary Table 1.**
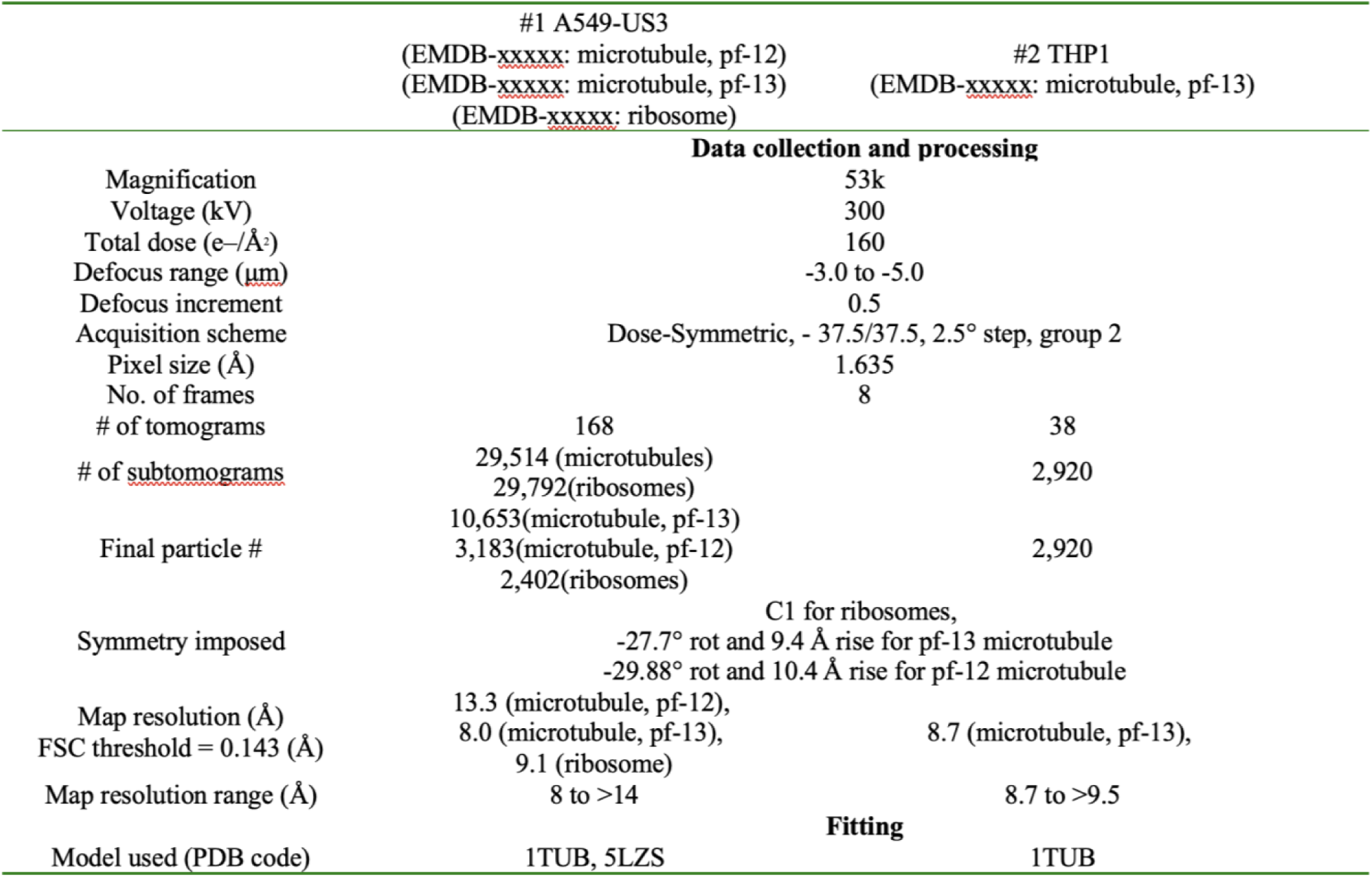
Cryo-ET data collection, processing, and map parameters.

**Supplementary Movie 1. Bidirectional lysosomal transport in MEFs.** Bidirectional transport of lysosomes through TNTs formed in primary MEFs stably expressing CTNS–GFP.

**Supplementary Movie 2. Bidirectional lysosomal transport in HeLa cells expressing US3–BFP.**

Lysosomal transport (LysoTracker) within TNTs formed in HeLa cells expressing US3–BFP kinase.

**Supplementary Movie 3. EB1–GFP dynamics in TNTs of HeLa cells expressing US3–BFP.**

**EB1–GFP comets visualized within TNTs in HeLa cells expressing US3–BFP.**

**Supplementary Movie 4. Bidirectional lysosomal transport in THP-1 cells treated with daunorubicin.**

Bidirectional transport of daunorubicin (magenta), internalized within lysosomes (LysoTracker), through TNTs.

